# A Comprehensive Human Embryogenesis Reference Tool using Single-Cell RNA-Sequencing Data

**DOI:** 10.1101/2021.05.07.442980

**Authors:** Cheng Zhao, Alvaro Plaza Reyes, John Paul Schell, Jere Weltner, Nicolás M. Ortega, Yi Zheng, Åsa K. Björklund, Laura Baqué-Vidal, Joonas Sokka, Ras Torokovic, Brian Cox, Janet Rossant, Jianping Fu, Sophie Petropoulos, Fredrik Lanner

**Affiliations:** Department of Clinical Science, Intervention and Technology, Karolinska Institutet, and Division of Obstetrics and Gynecology, Karolinska. Universitetssjukhuset, Stockholm, Sweden; Department of Integrative Pathophysiology and Therapy, Andalusian Molecular Biology and Regenerative Medicine Centre (CABIMER), Seville, Spain; Stem Cells and Metabolism Research Program, University of Helsinki, Haartmaninkatu 8, 00290 Helsinki, Finland; Folkhälsan Research Center, 00290 Helsinki, Finland; Department of Mechanical Engineering, University of Michigan, Ann Arbor, MI 48109, USA; Dept of Cell and Molecular Biology, National Bioinformatics Infrastructure Sweden, Science for Life Laboratory, Uppsala University, Husargatan 3, SE-752 37 Uppsala, Sweden; Department of Physiology, Faculty of Medicine, University of Toronto, M5S 1A8, Toronto, Canada; Program in Developmental and Stem Cell Biology, Hospital for Sick Children, Toronto, Ontario, Canada; Department of Cell & Developmental Biology, University of Michigan Medical School, Ann Arbor, MI 48109, USA; Department of Biomedical Engineering, University of Michigan, Ann Arbor, MI 48109, USA; Département de Médecine, Université de Montréal, Montréal Canada; Centre de Recherche du Centre Hospitalier de l’Université de Montréal, Axe Immunopathologie, H2X 19A Montréal Canada; Ming Wai Lau Center for Reparative Medicine, Stockholm Node, Karolinska Institutet, Stockholm, Sweden; Department of Biomedical and Chemical Engineering, Syracuse University, Syracuse, NY 13244, USA

**Keywords:** Single-cell RNA-sequencing, human development, embryo models

## Abstract

Stem cell-based embryo models offer unprecedented experimental tools for studying early human development. The usefulness of embryo models hinges on their molecular, cellular and structural fidelities to their *in vivo* counterparts. To authenticate human embryo models, single-cell RNA-sequencing has been utilised for unbiased transcriptional profiling. However, a well-organised and integrated human single-cell RNA-sequencing dataset, serving as a universal reference for benchmarking human embryo models, remains unavailable. Herein, we developed such a reference, through integration of six published human datasets covering developmental stages from the zygote to the gastrula. Lineage annotations are contrasted and validated with available human and non-human primate datasets. Using stabilised UMAP we constructed a web tool, where query datasets can be projected on the reference and annotated with predicted cell identities. Using this reference tool, we examined several recent human embryo models, highlighting the risk of misannotation when relevant references are lacking.

## Introduction

The use of stem cell-based embryo models mimicking human embryogenesis from the zygote stage to gastrulation has the transformative potential for advancing understanding of early human development. However, to establish the usefulness of these models, it is critical to validate and benchmark them against human embryos of corresponding developmental stages to ensure their resemblance and fidelity to the *in vivo* human embryos they aim to model. Such comparison and validation should be conducted at molecular, morphological, and when possible functional levels ^1,2^. Molecular characterizations of human embryo models are commonly conducted by examining expression levels of individual lineage markers. However, it is increasingly recognized that cell types and their states are not always distinguishable with individual or a limited number of markers, as many cell lineages in early human development share molecular markers. As such, global gene expression profiling becomes necessary and offers an opportunity for unbiased transcriptome comparison between human embryos models and their *in vitro* counterparts. Although still limited, individual human embryo transcriptome dataset references have been reported during the last 10 years, covering developmental stages from fertilisation to gastrulation ^3–8^. However, a well-organised and integrated human single-cell RNA-sequencing dataset, serving as a universal reference for benchmarking human embryo models, remains unavailable. Herein, we developed such a human embryogenesis reference dataset, through integration of six published human datasets covering developmental stages from the zygote to the gastrula. Lineage annotations are contrasted and validated with corresponding available human and non-human primate datasets. Based on this reference, we have performed comprehensive comparisons with recently reported human embryo models and further developed a robust, user-friendly online tool to explore the reference and benchmark stem cell-based embryo models.

## Results

### Development of human embryogenesis reference from the zygote to the gastrula

To create a human embryogenesis transcriptome reference covering developmental stages from the zygote to gastrulation, we collected six published datasets generated with single-cell RNA sequencing. We reprocessed these datasets, including mapping and feature counting, using the same genome reference (v.3.0.0, GRCh38) and annotation through a standardised processing pipeline (see Methods). This approach was adopted to minimise potential batch effects as much as possible. These datasets included cultured human preimplantation stage embryos, 3D cultured post-implantation blastocysts, and a Carnegie stage (CS) 7 human gastrula isolated *in vivo*, presumed to be day 16-19 ^3–8^ (Figure 1A). For integration we employed fast Mutual Nearest Neighbor (fastMNN) methods ^9^ to establish a high-resolution transcriptomic roadmap. In total, expression profiles of 3304 early human embryonic cells were embedded into the same 2-dimensional space (Figure 1A). Considering both the published and updated annotations, the resulting UMAP displays a continuous developmental progression with time and lineage formation (Figure 1B, C). The first lineage branch point occurs as the inner cell mass (ICM) and the trophectoderm (TE) cells diverge during embryonic day (E) 5 followed by the lineage bifurcation of the epiblast and hypoblast (Figure 1C, Figure S1E) ^3,7,10^. Early epiblast cells from E5 to E8 clustered together while the majority of the epiblast from E9 to CS7 formed a distinct cluster referred to as the ‘late epiblast’ cluster (Figure S1A-D). A similar transition was observed from early to late hypoblast, occurring around E10 (Figure S1A-D). Following extended 3D implantation culture the TE matures into Cytotrophoblast (CTB), Syncytiotrophoblast (STB) and Extravillous trophoblast (EVT) as described in Xiang et al., 2020 ^4^ (Figure 1B, C, Figure S1E). Within the gastrula we detect further specification of the epiblast into amnion, primitive streak (PriS), mesoderm and definitive endoderm (DE) together with extraembryonic lineages including yolk sac endoderm (YSE), extraembryonic mesoderm (ExE_Mes) and hematopoietic lineages (haemato-endothelial progenitors (HEP) and erythroblasts), in agreement with the original publication (Figure 1B, C, Figure S1E) ^5^.

**Figure 1:**
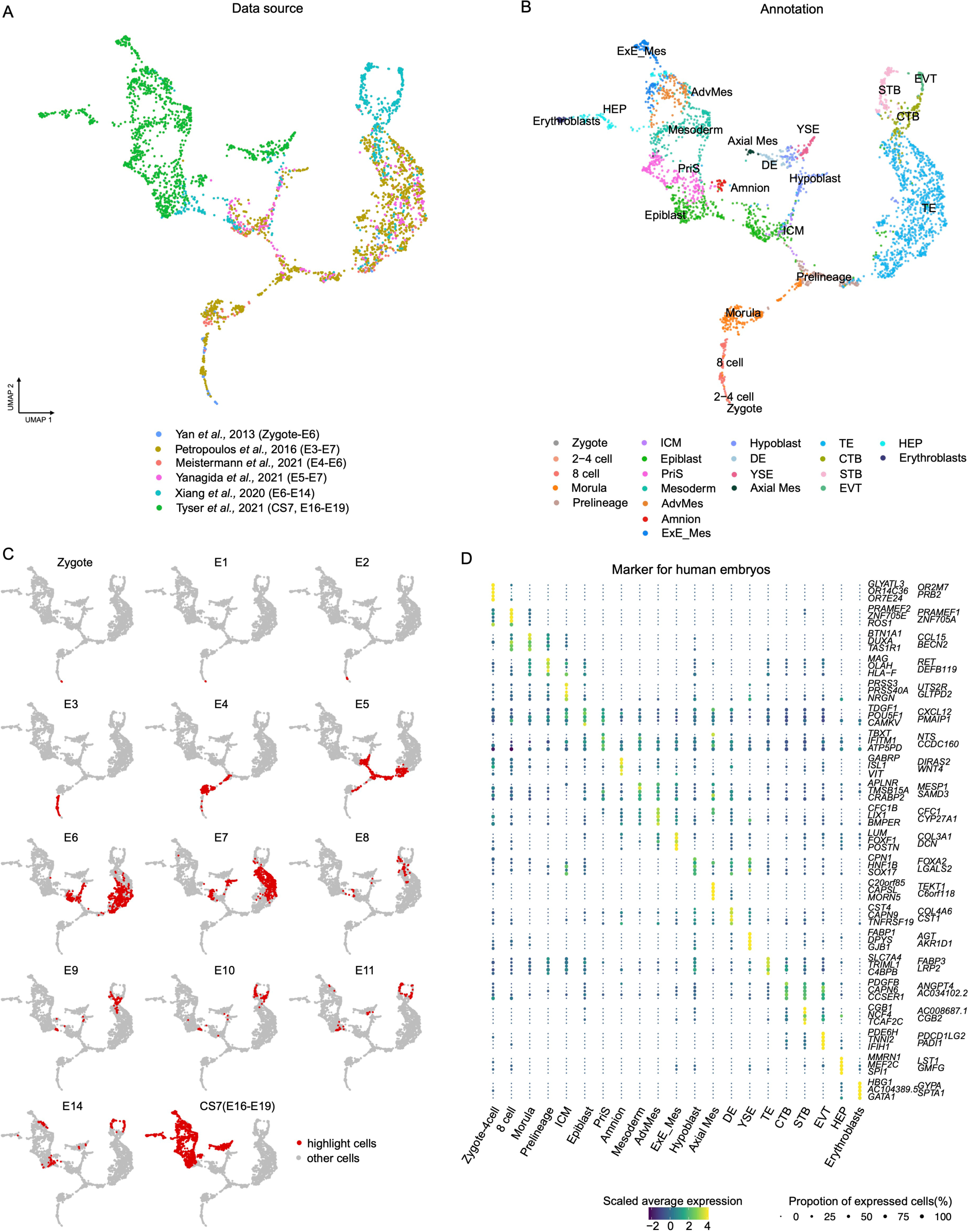
Construction of a human embryonic reference from zygote to the gastrula. (A) UMAP projection of the integration of six embryonic datasets. Colour of each data point represents the source of the data. (B) Similar to (A), but the colour indicates the cell annotations retrieved from each publication. (C) Highlight cells from different embryonic time points on the human embryonic reference. (D) Dot plot illustrating the expression of the top 5 lineage-specific genes used in the human embryonic reference. The size and colours of dots indicate the proportion of cells expressing the corresponding genes and scaled values of log-transformed expression, respectively.

Slingshot trajectory inference revealed three main trajectories starting from the zygote (Figure S1E). In the epiblast, hypoblast, and TE trajectories, 372, 345, and 253 transcription factor genes, respectively, were significantly associated with inferred pseudotime (Figure S2A, Table S1). Transcription factors such as *DUXA* and *FOXR1* exhibited high expression during morula stages but decreased expression across all three lineages (Figure S2A, C). Pluripotency markers like *NANOG* and *POU5F1* were expressed in the epiblast during preimplantation and decreased in expression following implantation whereas *HMGN3* showed increased expression at the later stages (Figure S2A, C). Along the hypoblast trajectory, *GATA4* and *SOX17* showed early expression while *FOXA2* and *HMGN3* demonstrated increased expression in the late phase. Within the TE trajectory, *CDX2* and *NR2F2* showed early expression while *GATA2*, *GATA3*, and *PPARG* showed increased expression during TE development to CTB (Figure S2A, C). *HMGN3* was also associated with the later stage of the TE trajectory in a similar manner as seen in epiblast and hypoblast trajectory (Figure S2A, Table S1), a pattern also observed in the non-human primate datasets ^11–15^. Comparing the epiblast with TE trajectory, genes like *ZSCAN10* and *NR2F2* were specifically associated with epiblast and TE trajectories, respectively, as they segregate from each other. Comparing epiblast with hypoblast trajectory, genes such as *GATA4* were specifically associated with the hypoblast trajectory (Figure S2B, C, Table S1). This data set provides an opportunity for further functional characterization of key transcription factors which may drive the differentiation of the three main lineages (Table S1).

Having assembled transcriptional profiles from zygote to the gastrula (CS7), we identified unique markers for each lineage throughout embryo development including the known expression of *DUXA* in morula ^16,17^ *PRSS3* in ICM cells ^10^, *TDGF1* and *POU5F1* in epiblast, TBXT in PriS cells, and *ISL1* and *GABRP* in amnion cells ^11,18^, *LUM* and *POSTN* in extraembryonic mesenchyme ^19^ (Figure 1D, Table S2). In addition, we identified genes such as *RBP4* ^20^ and *AFP* ^21^ that were specifically expressed in YSE but not in the hypoblast and DE (Figure S3A). When comparing the extraembryonic mesoderm with the embryonic mesoderm, genes including *DCN*, *ANXA1* and *POSTN* were specifically expressed in the ExE_Mes but not in embryonic mesoderm ^19^. In contrast, *ZNF738*, *TUBB2B*, and *NPY* were more enriched in embryonic mesoderm cells (Figure S3B).

As the human embryogenesis reference aligned well with reported annotations, embryo developmental timepoints, and marker expression, we also sought to investigate the key co-regulatory regulons in each lineage throughout embryo development. Therefore, we performed SCENIC (single-cell regulatory network inference and clustering) analysis ^22^ to explore the activities of transcription factors using MNN-corrected expression values across a wide range of embryonic time points. This analysis, captured known transcription factors corresponding to individual lineages, confirming the lineage identities. For example, we observed signatures of important transcription factors such as *DUXA* in 8-cell lineages ^7^, *VENTX* within the epiblast ^23^, *TEAD1* and *GATA2/3* within the TE, *TFAP2C* and *ISL1* in amnion ^11^, *E2F3* in erythroblasts, and *MESP1* in mesoderm ^23,24^ while extraembryonic mesenchyme is enriched in *HOXC8* signatures (Figure S1F).

Given the enrichment of key marker genes and transcription factor regulatory networks, we are confident that our embryonic reference provides reliable transcriptome profiles for each lineage present in early human embryo development included in those datasets.

### Integration with non-human primates

It should be acknowledged that the existing human embryo reference datasets are still limited with only a single *in vivo* gastrulating embryo ^5^. We therefore compared the six human embryo datasets with additional datasets from cynomolgus macaque, encompassing transcriptome profiles for preimplantation embryos ^13^, embryos collected during gastrulation ^14^, and *in vitro* cultured post-implantation embryos ^11^. Furthermore, two datasets for marmoset were included, comprising a transcriptome profile for preimplantation embryos ^12^ and implanted marmoset embryos at CS 5-7, including spatial information ^15^. Considering the effective performance of the MNN method in removing batch effects among datasets, similar MNN expression correction was performed within each species to eliminate intra-species batch differences. This was followed by Canonical Correlation Analysis (CCA) to simultaneously anchor the datasets ^25^. Through these efforts, single-cell transcriptional profiles from 11 datasets across three primate species were successfully embedded into the same 2-dimensional UMAP space (Figure 2A, B). The integrated UMAP revealed a continuum of transcriptional changes, which match the lineage annotations and developmental stages reported by the original publications (Figure 2A, Figure S4), and confirmed that intra/inter-species batch differences were effectively removed during the aforementioned process.

**Figure 2.**
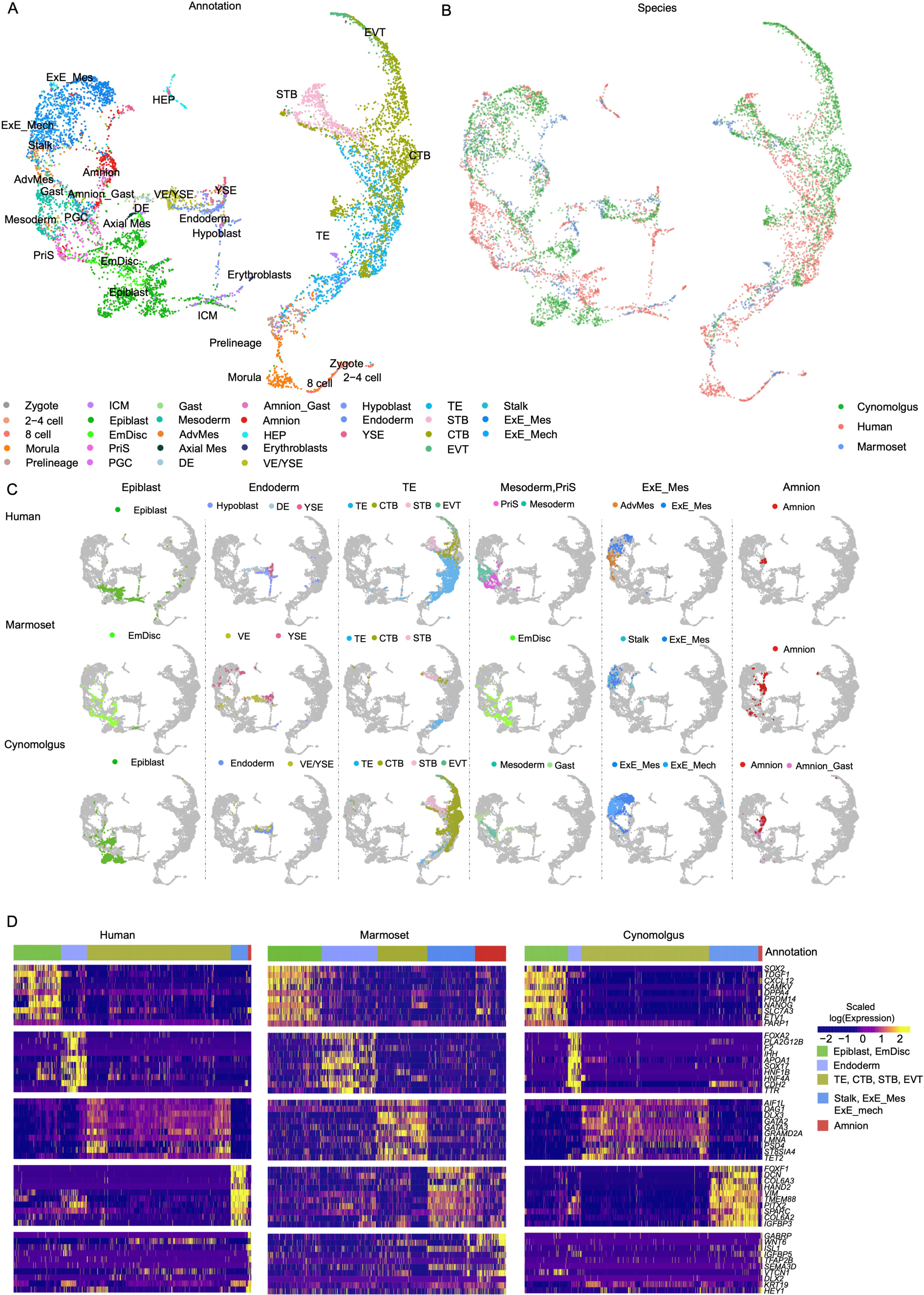
Cross-species integration involving cells from early human, cynomolgus monkey, and marmoset embryos. (A) UMAP projection of the integrated datasets from six human, three cynomolgus monkey, and two marmoset embryos. Each data point’s colour corresponds to the cell annotations retrieved from each publication. (B) Similar to (A), but the colour represents the species of the data. (C) Highlights cells from each lineage belonging to their respective species in the cross-species integration. (D) Expression of the top 10 lineage marker genes conserved in primate species.

From the cross-species integration, we observed that the prelineage to ICM and TE branch, ICM to epiblast or hypoblast branch, epiblast to PriS, mesoderm and amnion branch, and TE to CTB to STB or EVT branch were well conserved in all primate species (Figure 2A, C). Marmoset embryonic disc (EmDisc) aligned with epiblast-related lineages of the late CS7 human gastrula, including the ‘epiblast’, ‘primitive streak’, ‘mesoderm’, and E16-E17 epiblast cells in cynomolgus monkey (Figure 2A, C and Figure S4). Visceral (VE) and yolk-sac endoderm (YSE) aligned well between human and marmoset (Figure 2A, C). Marmoset cells reported with uncertain identity in the original report among CS7 secondary yolk sac (SYS), ExE_Mes, matched with SYS and amnion cells in our analysis ^15^. “Gastrula” (Gast) cells in cynomolgus monkey overlapped with human gastrula lineages, including the mesoderm and PriS cells. Some cells belonging to “Gast” cells mainly from E16 and E17 cynomolgus monkey (called “Late_Gast”) were aligned to part of the human advanced mesoderm (Figure 2A, C). The position of cells from “Amnion_Gast” from cynomolgus monkey ^14^ overlapped with cells from the human dataset, validating their identity as true amnion cells ^5^. In addition, we observed that extraembryonic cells (ExE_Mes cells, stalk, and extraembryonic mesenchyme (ExE_mech) formed large clusters. Human advanced mesoderm and cynomolgus “Late_Gast” cells, joined the embryonic mesoderm cells and extraembryonic cells, displaying transcriptional similarities shared by those late/advanced mesoderm cells and extraembryonic cells.

Utilising the comprehensive primate reference we identified conserved markers for the major lineages, including well-known pluripotency markers such as *SOX2* and *NANOG* in the epiblast, *SOX17* and *FOXA2* in endoderm cells, *GATA2* and *GATA3* in trophectoderm and its derivatives, *DCN* and *VIM* in extraembryonic cells, and *GABRP* and *ISL1* in amnion cells (Table S3) ^3,5,11,15,26,27^. In summary, we have compiled a comprehensive integration that encompasses single-cell RNA-sequencing datasets of early primate embryos spanning a wide range of time points. This provides a framework for investigating the concordances and variances among different species. Importantly, the extensive non-human primate datasets provide further support to the annotations in our human embryogenesis reference.

### Stabilised projection of query cells onto human embryonic reference

Having characterised the assembled embryonic reference, we next sought to establish a stabilised UMAP for evaluating the validity of stem cell-based embryo models. To achieve this, we distilled the entire fastMNN reference construction process into three major parts: 1) rescaling normalisation process, 2) PCA subspace projection after MNN correction, and 3) UMAP projection (Figure 3A). Throughout this analysis, adhering carefully to the assumptions that MNN pairs define the most similar cells of the same type across batches and that batch effects should be nearly orthogonal to the biological subspace ^9^, we divided the query data into small samples and filtered ambiguous pairs (see in Methods). After generating comparable normalised data for the query dataset, it was projected onto the same PCA subspace and subsequently corrected, followed by projection onto the same UMAP space as the reference (Figure 3A). After the query cells were successfully projected onto the reference, cell identities were assigned based on the stabilised UMAP coordinates. Query cells which do not correspond to those in our reference, were filtered out based on correlation filtering and annotated as “non-related” (see Methods).

**Figure 3.**
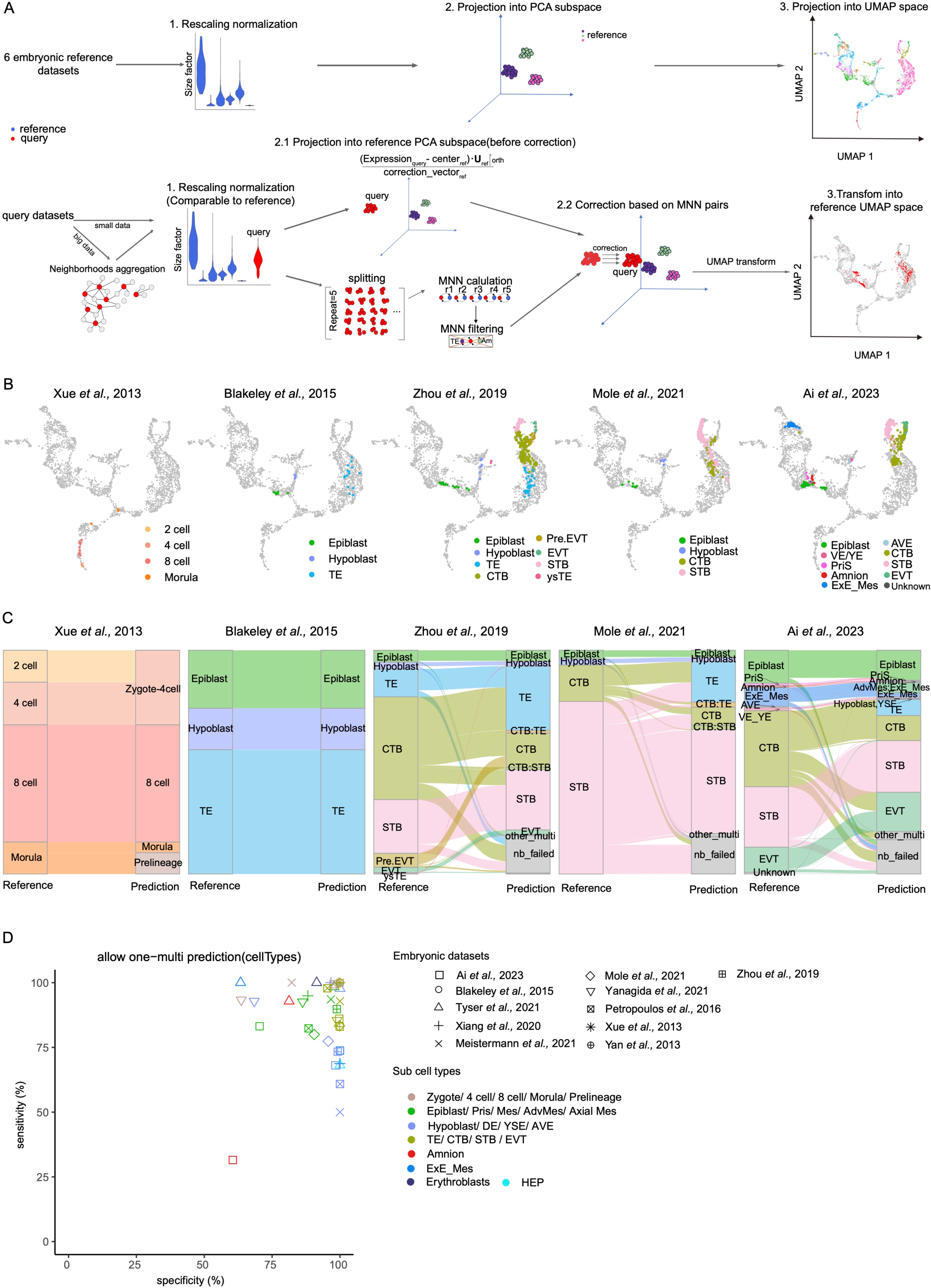
Validation of early embryogenesis prediction tool. (A) Workflow for processing projection of query cells onto the human embryonic reference. First thequery data underwent rescaled normalisation to ensure that expression values were comparable to those of the reference dataset. This step was taken after deciding whether to aggregate cells into neighbourhoods for low-depth, large datasets. Cosine normalisation of query expression involved removing the same grand centre values from reference calculations and performing a dot product calculation with a unitary matrix obtained from singular value decomposition during reference construction. Subsequently, orthogonalization was applied to remove variation along the reference batch correction vector, projecting it onto the same PCA subspace as the reference. Simultaneously, the query dataset with more than 200 cells or neighbourhood nodes was divided into small samples, repeated five times, and fed as input to calculate MNN pairs with the reference datasets separately. Uncertain MNN pairs, assigned to TE and Amnion simultaneously, were then removed. Using the filtered MNN pairs, a batch correction vector was computed to correct the PCA coordination of the query dataset. Finally, the UMAP embedding was transformed using the UMAP model calculated from the reference dataset. (B) Projection of cells (or neighbourhood nodes) onto the human embryonic reference from four other published embryonic datasets. The colour of each data point represents the cell annotations retrieved or restored for each publication. Light grey data points represent cells used in embryonic reference construction. (C) Alluvial plot comparing the original cell-type annotations to the predicted identities obtained from the early embryogenesis prediction tool. Predictions identified as “ambiguous” or “nb_failed” represent cells with uncertain predictions or cells that fail to form neighbourhoods, respectively. (D) Prediction sensitivity and specificity for each cell type in the embryonic datasets. The shape and colour of data points indicate queried cell types and data sources, respectively.

To test the prediction performance, five additional embryonic datasets were utilised, including a pre-lineage embryo dataset spanning from the 2-cell to morula stage ^28^, a blastocyst dataset ^27^, and three peri-implantation and post-implantation dataset ^29–31^ (Table S4). Of note, regardless of the library preparation utilised to generate these datasets (e.g. SmartSeq2, Trio-seq, 10X sequencing) or whether the datasets were used as processed by the authors or reprocessed in-house, all five datasets displayed excellent alignment with their counterparts from our embryonic reference (Figure 3B, C and Figure S5A-C). Overall, we achieved 87.6% sensitivity and 92.9% specificity, when comparing published annotation of the main lineages during embryo development with our prediction (Figure 3D). When considering a more granular classification of sub-lineages, we obtained a 76.0% sensitivity / 88.3% specificity and 80.5% sensitivity / 86.8% specificity, respectively, depending on the stringency set to allow multiple predictions for one cell (Figure S5D). In general, this predictive pipeline delivers a highly accurate performance for scRNA-seq datasets, irrespective of upstream library preparation or data processing, which can be leveraged for benchmarking embryonic cells with embryonic models.

### Mapping stem cells and derived lineages

Naïve and primed human pluripotent stem cells (hPSCs) are considered analogous to the embryonic epiblast at pre- and post-implantation, respectively. As expected, naïve cells mapped to the early epiblast cells before E9, whereas primed cells were projected to the late epiblast cells (Figure 1C, Figure S1C and Figure 4A). Trophoblast stem cells, derived from blastoids using Okae et al., 2018 culture conditions ^32,33^, mapped to E7-E10 TE and CTB cells, confirming a trophoblast identity. There is a longstanding debate whether human naïve and primed stem cells can be converted to trophoblast lineage ^2,34,35^. One concern has been that converted putative trophoblast cells instead may be misannotated ExE_Mes or amnion lineage as some lineage markers are shared ^36,37^. For this reason, we assessed the transcriptional profiles of published TE-like cells (TLC) derived from naïve and primed hPSC in addition to our own unpublished data-set of primed derived TLC (In-house) (Figure 4B) ^34,38–41^. Both naïve and primed derived cells were predicted as TE cells of the blastocyst or later trophoblast stages. However, primed derived cells also included significant contributions of both amnion and ExE_Mes cells (Figure 4C). These predictions were confirmed by module score calculation based on lineage marker genes (Figure S6A and B). Given the different represented epiblast stages for naïve and primed, among the 705 differentially expressed genes (DEGs) between naïve and primed cells, 39, 47, and 48 were also differentially expressed in the alternative lineages amnion, ExE_Mes, and PriS compared with preimplantation TE cells (Table S5). Interestingly, the DEGs shared by the alternative lineages were also expressed specifically in the primed cells (17 out of 17 genes). Conversely, genes preferentially expressed in the TE cells were also enriched in naïve cells (53 out of 60 genes) (Figure S7 and Table S5). The genes shared by naïve and TE cells, including the key transcriptional factors *NLRP2*, *NLRP7*, *TFAP2C*, *KLF4*, and *ELF3* may facilitate naïve cells to adopt a TE fate over alternative fates including amnion, ExE_Mes and PriS (Table S5). We next compared the TLCs generated from both stem cell states, with the embryonic preimplantation TEs. We identified 50 and 71 DEGs consistently differentially expressed in naïve or primed TLCs when compared with TE cells (Figure 4D and Table S6). 30 of them shared in both naïve and primed derived TLCs (Figure 4E and Table S6). TE transcription factors such *CDX2* and *GTSF1* were expressed at low levels in primed-derived cells, while *HES4* (hes family bHLH transcription factor 4), which participates in transcriptional regulation and the Notch signalling pathway, was expressed only in stem cells-derived TLCs. *DNMT3L* involved in the process of DNA methylation was expressed at low levels in all stem cells, and was particularly low in primed-derived cells (Figure S8). *XIST* levels were also absent in primed derived TLC while some naïve-derived TLC expressed significant levels (Figure 4E, S7) suggesting that primed derived TLC might have an aberrant epigenetic state.

**Figure 4.**
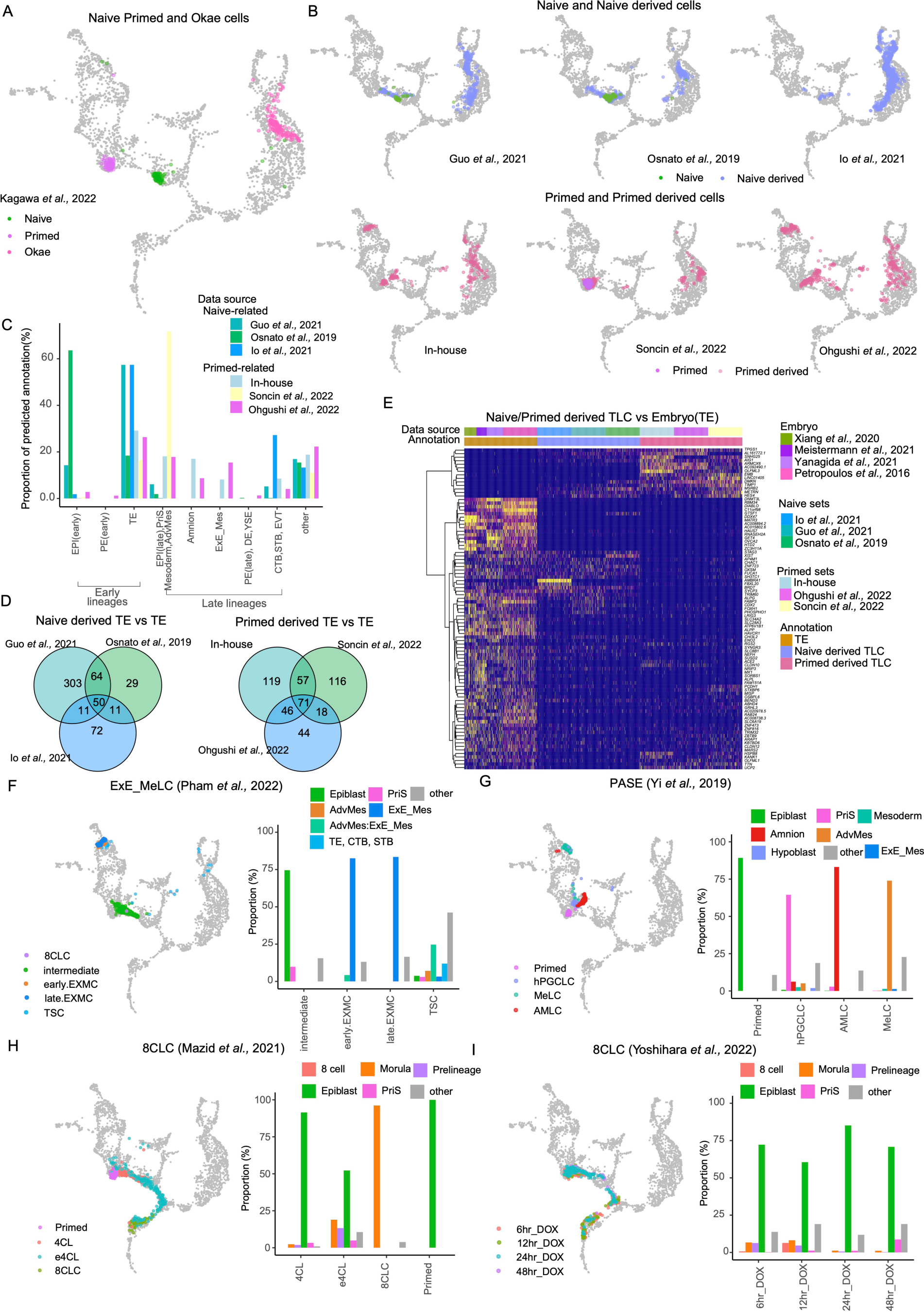
Application of early embryogenesis prediction tool on stem cell models. (A) Projection of naïve, primed, and Okae cells from Kagawa et al., 2022 onto the human embryonic reference. The colour of each data point represents the cell annotations retrieved from original publication. Light grey-coloured data points represent cells used in embryonic reference construction. (B) Projection of six datasets that use naïve or primed cells to model TLCss. The colour of each data point represents whether the cells (neighbourhood nodes) are naïve, primed, or their derived cells. (C) Barplot showing the proportion of predicted cell identities for naïve-derived and primed-derived cells. (D) Venn diagram showing the overlap of DEGs between naïve-derived preimplantation TLC and embryonic preimplantation TE cells, as well as the overlaps of DEGs between primed-derived preimplantation TLCs and embryonic preimplantation TE cells. (E) Heatmap showing the expression of DEGs in preimplantation TLCs and embryonic preimplantation TE cells. The DEGs were conserved in all three naïve-derived TLCs comparisons or conserved in all three primed-derived TLCs comparisons, primed-derived TLCs, and embryonic TE cells. (F, G) Projection of cells (neighbourhood nodes) from two studies modelling 8-cell-like cells. The colour of each data point represents the cell types or timepoint retrieved or restored for each publication. Light grey-coloured data points represent cells used in embryonic reference construction. Barplot showing the proportion of predicted cell identities stratified by cell types or timepoint. (H, I) Projection of cells (neighbourhood nodes) from two studies modelling extraembryonic mesoderm cells and post-implantation amnion sac (PASE). The colour of each data point represents the cell annotation retrieved or restored for each publication. Light grey-coloured data points represent cells used in embryonic reference construction. Barplot showing the proportion of predicted cell identities stratified by cell original annotation.

We next wanted to apply our reference map to evaluate published examples of converting stem cells not into TLC but specifically into ExE_Mes and amnion ^19,42^ and confirmed the accurate annotation of naïve derived ExE_Mes cells as well as amnion in the stem cell-based post-implantation amniotic sac embryoid (PASE) model (Figure 4F, G). The PASE model also includes primitive streak and mesoderm-like cells which map as expected while Primordial Germ Cell-Like Cells (hPGCLC) map to the primitive streak. This highlights the point that our current reference can not resolve PGCs as the original dataset only included 7 annotated PGC which was not enough to establish a discrete cluster on our reference UMAP.

Another exciting development is transient conversion of embryonic stem cells into 8-cell like cells (8CLC) states. Indeed, we observed a large proportion of morula-like cells but not 8CLC in Maizd et al., 2022, confirmed by module score calculation (Figure 4H and Figure S6C) ^43^. In Yoshihara et al., 2022 we observed around 0.7% and 6.4% 8CLC at 6 hours and 12 hours after transient doxycycline treatment in Doxycycline-inducible DUX4-TetOn hESCs, respectively, which aligned with the reported 6.6% of DUX4-pulsed cells which were converted to a state that transcriptionally resembled 8-cell stage embryo 12 h after induction (Figure 4I) ^44^.

### Evaluating the preimplantation blastoids models

Next we explored the transcriptional profiles of blastoids established from naïve and EPSCs or through partial reprogramming ^6,32,45–49^. Projection of three naïve hPSC-derived blastoids reveals the presence of expected cell types, with the majority of annotated epiblast-like cells (ELC), hypoblast-like cells (HLC), and TLC overlapping with their counterparts in the human blastocyst embryo although a fraction showed signatures more in line with postimplantation stages (Figure 5A, B, and Figure S11B). In addition, small fractions of cells with signatures resembling ExE_Mes and amnion were also detected (Figure 5A). In blastoids generated from reprogrammed stem cells ^46^, the majority of TLCs overlap with the amnion reference (Figure 5B and Figure S9). Blastoids cells derived from EPSCs ^48^ were reported to have the morphology of a blastocyst but lack correct transcriptional profiles. In agreement with that finding, most cells were predicted as ExE_Mes or advanced mesoderm (AdvMes). EPSC-derived blastoids Fan et al., 2021 ^47^ did contain TE-like cells but the majority of the cells resembled “late epiblast” cells (after E10) with very few HLCs (Figure 5B). To validate these potential issues in an independent manner, we utilised published cynomolgus data ^11^, which includes both amnion and trophoblast cells within the same dataset. Comparative transcriptome analysis of blastoids using cynomolgus data as a reference clearly shows that the majority of TLCs in blastoids of Liu et al., 2021 (Figure S9A) cluster with the amnion rather than the TE lineage (Figure S9A). The majority of TLCs in blastoids from Sozen et al., 2021 (Figure S9A) are closely related to ExE_Mech. HLCs from Fan et al., 2021 did not align with the reference endoderm cells. Additionally, TLCs in blastoids from Liu et al., 2021 lack or express low levels of TE markers like *GATA2*, *GCM1*, and *BIN2* but express amnion markers *ISL1*, *GABRP*, and *IGFBP7* (Figure S9B, C). Furthermore, we utilised DEGs between early and late stages of the epiblast and hypoblast cells (Figure S1D and Table S7) and checked their expression in blastoids (Figure S10). Original ELCs of blastoids from EPSC ^47,48^ preferentially express late epiblast DEGs but lowly expressed early epiblast DEGs, similar to primed cells (Figure S10A). HLC of blastoids from reprogrammed cells ^46^ and EPSC cells ^47,48^ expressed DEGs enriched in late hypoblast cells. Predicted annotations were further justified by module score calculation (Figure S11). Overall, these results confirm results following the analysis using the embryogenesis reference tool. Our analysis supports that naïve derived blastoids are composed of cells with transcriptional profiles in line with blastocysts. However, detailed analysis still identifies 4, 7, and 22 DEGs shared by all three naive blastoids when compared with the human reference (Figure 5D, E). *PEBP1* and *TNNT1* were ectopically expressed in ELC, *DPP4* and *HAVCR1* were lacking in HLC and *ACE2*, *FMR1NB*, and *SLC6A19* were lacking in TLC of blastoids. It is not clear if these DEGs translate into functional differences but such aberrant expressions should be taken into consideration when using these models. One recent example is modelling of SARS-CoV-2 placenta infection using stem cell models. SARS-CoV-2 infects cells via its spike (S) protein binding to the host entry receptor *ACE2* (Angiotensin-converting enzyme 2). Several studies have examined SARS-CoV-2 infection and *ACE2* expression in trophoblast organoids and in alignment with our analysis of blastoids, they also have seen reduction of *ACE2* levels and SARS-CoV-2 infection in such *in vitro* models, highlighting important functional differences between stem cell models and embryonic cells ^50,51^.

**Figure 5.**
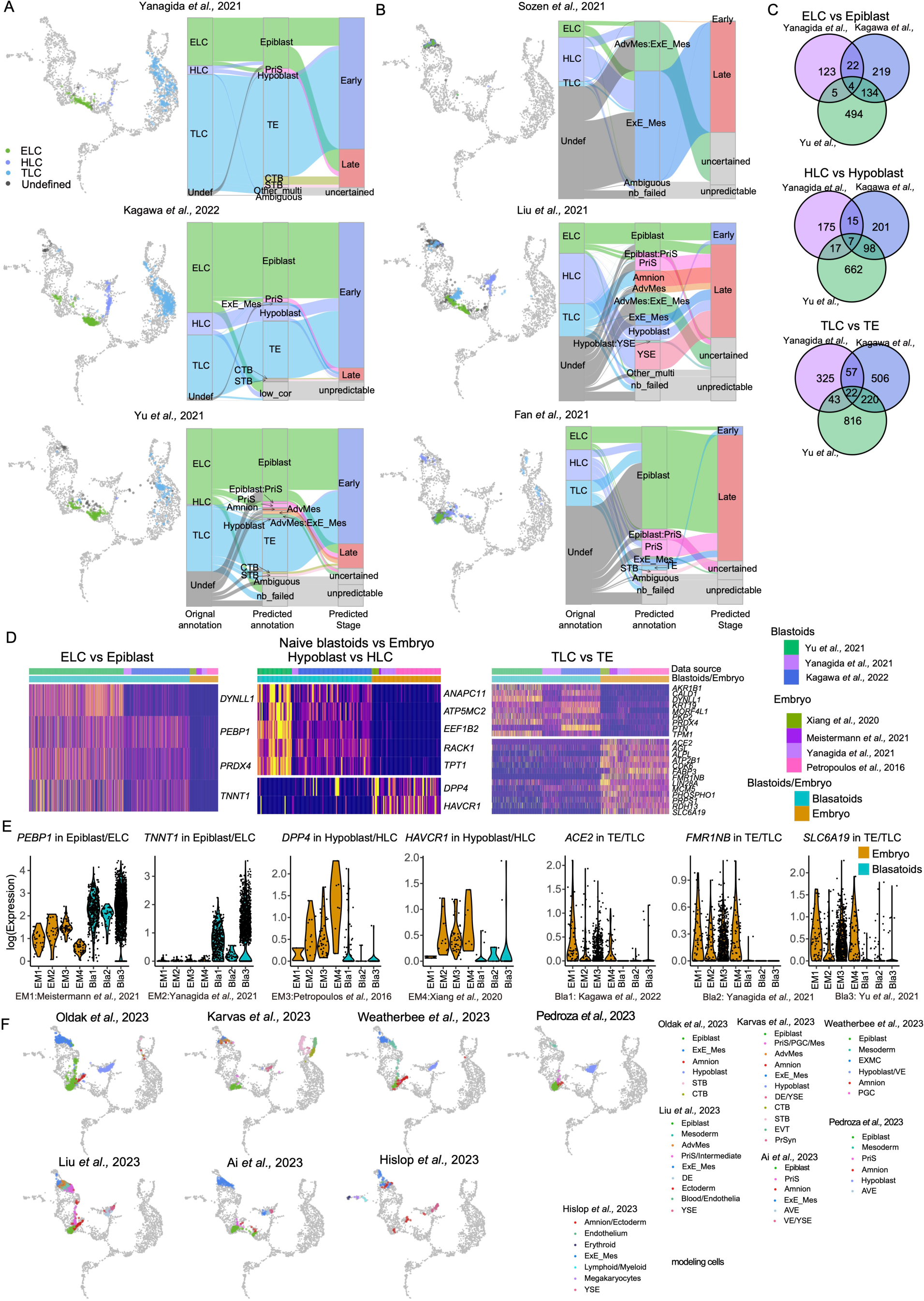
Application of early embryogenesis prediction tools on pre-implantation blastoid models. (A, B) Projection of blastoid cells (or neighbourhood nodes) onto the human embryonic reference. The colour of each data point represents the cell annotations retrieved or restored for each publication. Light grey-coloured data points indicate cells used in embryonic reference construction. Alluvial plot comparing original cell-type annotations (epiblast-like cells (ELC), hypoblast-like cells (HLC), and TLC from the six blastoids) to the predicted identities obtained from the early embryogenesis prediction tool. Naive-derived blastoids in (A) and reprogrammed or EPSC derived blastoids in (B). (C) Venn diagram showing the overlaps of DEGs between blastoids with pre-implantation embryonic lineages for three naïve-derived blastoids. (D) Heatmap showing the expression of conserved DEGs in all three naïve-derived blastoids between blastoids mimic lineages and embryonic reference. For visualisation, cells from the same cell type were randomly down-sampled to 200. (E) Violin plot showing the expression of representative DEGs between blastoids and embryonic reference. (F) Projection of post-implantation blastoid cells (or neighbourhood nodes) on human embryonic reference. The colour of each data point represents the cell timepoint retrieved for each publication. Light grey-coloured data points indicate cells used in embryonic reference construction.

### Evaluating stem cell derived postimplantation embryo models

Following implantation, the human embryo undergoes significant organisational changes crucial for gastrulation and subsequent development. Several post-implantation stem cell-derived models have been developed to mimic the development of human embryos during this stage. In addition to capturing morphogenesis and structural similarities to human post-implantation embryos, a crucial consideration is whether these models encompass all key lineages of the developing early post-implantation embryo. To address this, we projected recent studies ^29,51–56^ onto our human embryogenesis reference to evaluate their similarities with post-implantation lineages (Figure 5F). Three studies combine pluripotent cells with hypoblast-induced cells, generating cells with signatures aligning with the embryogenesis reference cells including epiblast, PriS, mesoderm, ExE_Mes, amnion, and late hypoblast ^29,53,54^ (Figure 5F). Two other studies also included TLC in the assembloids, cultured in suspension ^52,55^, while the third started from naïve-derived blastoids which were allowed to grow attached in 2D or imbedded in 3D ^51^. The blastoid based model had the best contribution to the TE compartment in the embryonic reference (Figure 5F), possibly due to the attachment strategy which might better support TE expansion. It is important to note that postimplantation models do not necessarily need to capture all lineages of the human embryo to be useful. This is exemplified by the recent study by Hisplo et al. 2023 ^56^ which modelled postimplantation development without a TE compartment but succeeded in establishing yolk sac hematopoiesis which was evident also from mapping to the embryogenesis reference (Figure 5F)^56^. Our tool did not resolve amnion cells from Pedroza et al. and Karvas et al., possibly due to two distinct reasons. For Pedroza et al., their amnion-annotated cells are projected together with epiblast and PriS in our tool. Using the published annotations and gene expression matrix from Pedroza et al., we observed that only 62 out of 460 amnion-annotated cells exhibited at least one read for any one of the amnion markers, *ISL1, GABRP*, *VIT*, *VTCN1*, *WNT4*, or *WNT6*. It is still possible that there are early amnion cells, which are lacking in the Tyser et al, reference. For the Karvas et al, the amnion cells are less than 15 and interspersed with AdvMes-like cells in the original report ^51^. In this case the amnion cells are likely too few to form an independent amnion neighbourhood, but instead mix with ExE_Mes cells in our reference map.

### A web platform for embryonic study

During this work we have curated a total of 38 processed datasets, including their raw expression data, original annotation, and predicted annotation. Among these datasets, eight human embryonic datasets, six preimplantation blastoids datasets, one 8-cell-like-cell dataset, and one PASE dataset were reprocessed by us using the same mapping strategy (10X using cellranger ^57^ and non-10X using STAR ^58^) and genome annotation ^3–8,27,29,32,44–48^. This effort has allowed us to eliminate batch differences arising from different processing pipelines or gene annotations, providing a valuable resource for future investigations.

To facilitate comparative studies and easy access to analysis of single cell datasets related to the developmental stages of zygote to gastrulation, we have developed an interactive online tool based on the human embryogenesis reference and the extended primate cross-species embryonic reference using ShinyCell ^59^ (Figure 6A). This can be used to browse the expression of selected genes in both the human and primate reference atlas. Furthermore, we have transformed the prediction pipeline into an online analytical application which we entitled the Early Embryogenesis Prediction Tool (Figure 6B). Our primary focus was to create a user-friendly interface, ensuring that researchers can directly access and utilise it by providing only the gene expression matrix. The entire process of normalisation, projection, and annotation takes less than 10 minutes for less than 7000 cells (Figure 6C). We envision broad usage of these assembled datasets and online tools for the authentication of human embryo models, and in turn supporting the development and application of this emerging field.

**Figure 6.**
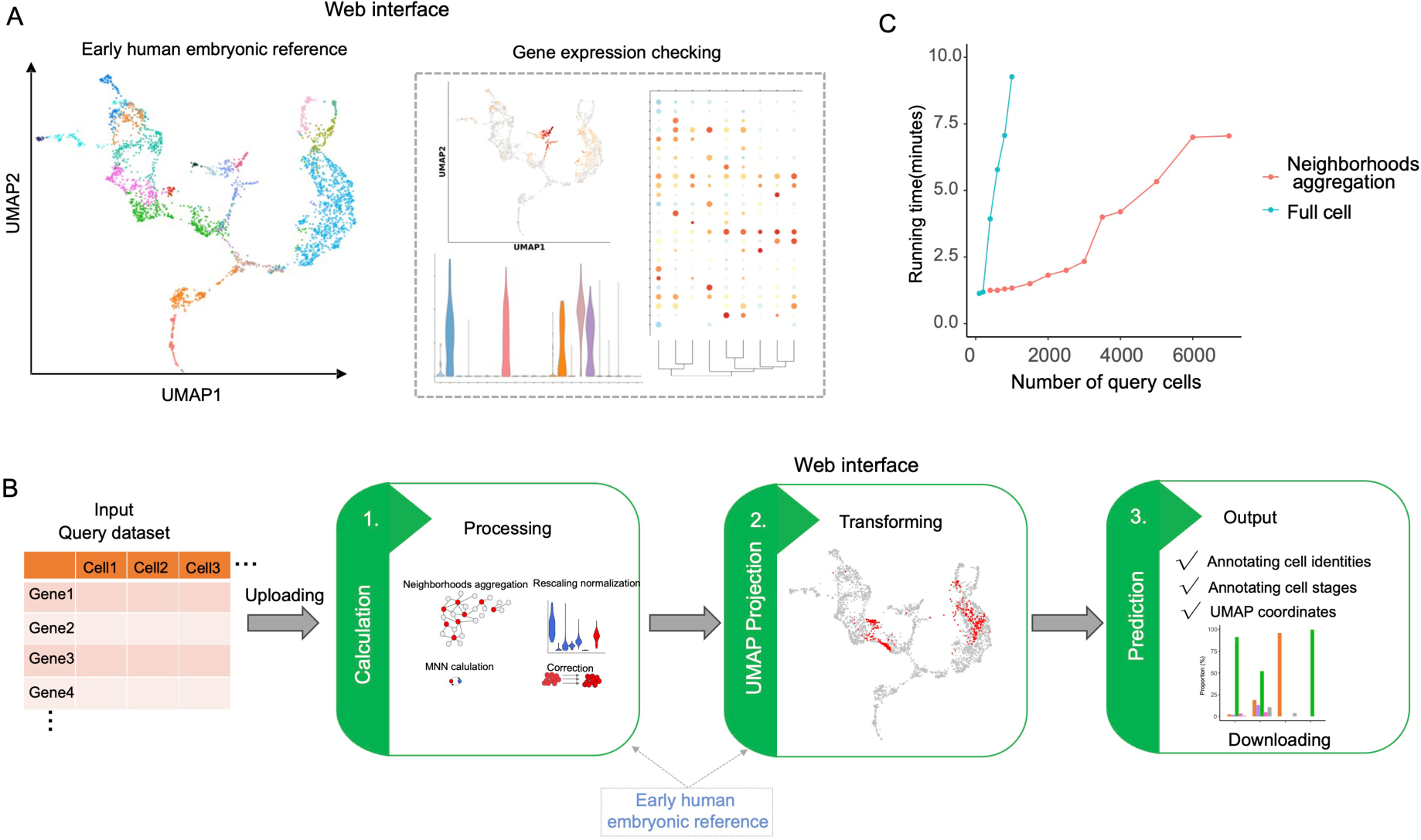
Web interface for our online resources. (A) Schematic of the web interface for the human embryonic reference. (B) Schematic of the web interface for the early embryogenesis prediction tool. (C) Running time for webtool with different numbers of query cells.

## Discussion

In this study, we have generated a comprehensive human embryo reference map by successfully integrating six embryonic datasets spanning from the zygote to the gastrulation stage. We identified markers encompassing all lineages during early embryo development, providing distinct clusters that resolved different lineages at various developmental stages. Leveraging the reference map we further interrogate published stem cell based models of human embryogenesis. This analysis highlights the risk of mis-antotating lineages of the early human embryo if relevant references are not included. It is particularly difficult to distinguish TE and amnion signatures as they share many markers. Using our reference UMAP, we identified three main trajectories (stemming from the EPI, PE or TE) and identified key transcriptional factors associated with their respective developmental trajectories. We then examined regulons comprising transcriptional factors and corresponding target genes for each lineage, to identify key transcriptional factors which may be driving the differentiation of these lineages. While several key transcriptional factors were identified, their specific functional roles require further investigation.

By distilling the MNN correction into three parts, we developed a pipeline that generates accurate stabilised UMAP projections with reference datasets when querying new datasets. Adhering to the requirements of MNN correction and considering the uniqueness of human embryonic reference datasets, we validated the pipeline with additional human embryonic datasets. The pipeline effectively distinguished early human embryo identities from different stages and lineages, displaying a specificity and sensitivity of 92.9% and 87.6%, respectively.

There are important limitations of our tool that should be considered when interpreting results obtained. First, an inherited limitation of MNN correction makes it challenging to correct datasets without shared similar identities. False-positive predictions may therefore occur for cells with transcriptome signatures which are not present in our reference but share close similarities. We partially resolved this issue by including correlation filtering in our processing pipeline. To test this we used unrelated datasets, including five public scRNA-seq datasets from human pancreas, macrophages, fetal kidney development and liver tissues (see Methods) and determined that a median top correlation coefficient threshold of 0.5 can eliminate unrelated cell types. However, this is still an important aspect to bear in mind. Another limitation is that the embryonic reference dataset contains few PGCs, with similarities in gene expression to primitive streak, and could therefore not be resolved as a distinct cluster which means that we can not identify PGCs using our tool. This could be resolved in the future with added in vivo reference datasets. Distinguishing between ExE_Mes cells and embryonic mesoderm cells proved challenging due to shared gene expression similarities and lack of well defined markers. Cross-referencing with the marmoset dataset with spatial information allowed us to reclassify some Adv_Mes cells as ExE_Mes cells. Genes specifically expressed in extraembryonic mesoderm cells, such as *DCN*, *ANXA1*, and *POSTN* ^19^, may be helpful in addressing these issues. Nonetheless, clear boundaries between these cell types remains a challenge and deeper understanding about these lineages are needed.

miloR is a tool designed for complex single-cell datasets which we utilise to form neighbourhoods in order to aggregate gene expression of cells. Our previous analyses ^55^ confirmed that neighbourhoods were approximately homogeneous in their cell-type composition, providing improved prediction accuracy by addressing sparsity in 10X-type datasets through gene expression aggregation within the same neighbourhood. While this aggregation method is beneficial for analyses with sparse datasets, it should be noted that a certain proportion of cells may fail to form neighbourhoods (labelled as “nb_failed” in prediction), leading to information loss for those cell types. Furthermore, if a query dataset contains a very rare subpopulation with limited number of cells, our tool may be unable to form independent neighbourhoods of that rare subpopulation and instead merge those cells with the closest resembling cells. Indeed, this was observed with the few amnion cells in the Karvas et al., dataset.

Our study has also provided insights into the complexity of stem cell states by comparing the transcriptional profiles of naïve and primed human pluripotent stem cells (hPSCs) to early embryonic lineages. The mapping of naïve cells to the early epiblast and primed cells to the late epiblast stages resonates with their developmental potential and distinct molecular characteristics. This alignment supports the notion that naïve and primed hPSCs mimic distinct developmental time points. The longstanding debate over the ability of human naïve and primed stem cells to transition into trophoblast lineages has also been addressed in our study, presenting a nuanced view of this complex process. The identification of trophoblast stem cells cultured under specific conditions mapping to the expected embryonic stages reinforces the notion that, with precise environmental cues, both primed and naïve stem cells can be guided into specific lineage pathways. However, the significant presence of amnion and extraembryonic mesodermal cells in primed-derived cell populations underscores the intricate balance and potential for diverse differentiation outcomes inherent in these cells. This finding underscores the necessity for refined cellular markers and rigorous validation strategies to ensure accurate lineage specification. Although both naive and primed stem cells can make TLCs, the transcriptional profiles still differ from embryonic equivalents. Our analysis identifies the same concern for current blastoids which at best do contain the expected cell types, but where significant transcriptional differences still exist. From the primed derived TLCs it appears that part of the problem is related to epigenetic dysregulation as these cells show reduced levels of DNA (cytosine-5)-methyltransferase 3-like (DNMT3L) and long non-coding RNA XIST which mediated X-chromosome silencing. In this aspect, the naive derived trophoblast-like cells are more alike the embryo trophoblast lineage. It will be interesting to see if starting from an even earlier stem cell state could give added value to embryo models. From that perspective it is exciting to see that transient conversion to earlier states, such as the 8-cell or morula-like stages could be confirmed in our embryogenesis tool. We hope this tool will facilitate development of more refined systems to study early human embryogenesis and benchmark these against the human embryo.

## Methods

### Data Availability

Datasets utilised in this study were obtained as summarised in Table S4. These include eleven human embryonic datasets covering various stages ^3–8,27–31^. Additionally, we included seven preimplantation blastoid models ^6,32,45–49^, seven postimplantation stem cell based models ^29,52–56,60^, three studies involving naïve cells giving rise to trophectoderm (TE)-like cells ^34,38,41^, three studies involving primed cells giving rise to TE-like cells (one generated in our lab) ^39,40^, two studies analyzing 8-cell-like cells ^43,44^, one study including post-implantation amniotic sac embryoid (PASE)-model ^42^, and one study mimicking extraembryonic mesoderm like cells ^19^. Furthermore, we included two embryonic datasets from Callithrix jacchus (Marmoset) ^12,15^ and three embryonic datasets from Macaca fascicularis (Crab-eating macaque) ^11,13,14^.

### Differentiation of primed derived Trophoblast like Cells

Primed hPSC growing on hrLN-521 (10 μg/mL, Biolamina, LN521-02) and NutriStem hPSC XF (Biological Industries, 05-100-1A) were enzymatically dissociated and seeded onto new plates also coated with hrLN-521 at a cell density of 1,84×104 cells/cm^2^. 24h after seeding, hPSC were moved to a 5% CO^2^/5% O^2^ incubator and were differentiated into trophoblastic cells using NutriStem hPSC XF without bFGF and TGFb that was supplemented with 10ng/mL BMP4 (R&D Systems, 314-BP-050e), 1µM A83-01 (R&D Systems, 2939) and 0,1µM PD173074 (Sigma-Aldrich, P2499) (BAP treated) according to a previously published protocol ^61^. The medium was changed daily during the 4 days of differentiation.

### In-house library preparation for scRNA-seq

Cells were dissociated with TrypLE Select (4min at 37°C; ThermoFisher Scientific, 12563011) and deactivated by adding 3 times more volume of NutriStem hPSC XF. Cells were collected in 0.04% BSA (Sigma-Aldrich, A7979) in PBS-/- and used following the Cell Multiplexing Oligo (CMO) Labeling for Single Cell RNA Sequencing Protocols with Feature Barcode technology protocol (10x Genomics, CG000391). Following cell count and verification of live cells using the NC-200 Nucleocounter (Chemometec) of individual samples and the final pool (4 CMO samples combined, aiming for 2000 cells per sample), we then used the Chromium Next GEM Single Cell 3′ Reagent Kits v3.1 (Dual Index) with Feature Barcode technology for Cell Multiplexing (10x Genomics, CG000388) according to the recommended user guide. CMO libraries and transcriptome libraries were sequenced using Illumia NextSeq 2000.

### Pre-processing of human scRNA-seq data and gene expression quantification

In-house 10X Genomics multiplexed primed-derived trophoblast scRNA-seq data were processed using “cellranger multi” pipeline (v.6.1.1) with default parameters ^57^. Published scRNA-seq data were processed using the “cellranger count” pipeline (v3.0.0). The STAR aligner (v2.5.1b) ^58^ was employed to map reads to the GRCh38 reference genome (v.3.0.0, GRCh38, downloaded from the 10X Genomics website). To minimize differences associated with the sequencing platform and data processing, published Smart-Seq2 datasets were also remapped to the same reference using the same aligner with default settings. Only uniquely mapped reads were retained for gene expression quantification. Raw read counts were further estimated using rsem-calculate-expression from the RSEM tool, with the option of “--single-cell-prior” ^62^. Datasets without a note indicating “reprocessed” in Table S4 were based on the previously processed expression matrix reported in their original publication.

### Pre-processing of scRNA-seq data and gene expression quantification for Marmoset monkeys (Callithrix jacchus) and Cynomolgus macaques (Macaca fascicularis)

The scRNA-seq transcriptomes of Marmoset embryos were sourced from studies in Boroviak et al., 2018, and Bergmann et al., 2022 ^12,15^. Expression matrices from both projects were obtained from https://github.com/Boroviak-Lab/SpatialModelling. Cells derived from maternal material were excluded from the entire analysis. For Cynomolgus macaques, embryonic scRNA-seq transcriptomes were downloaded from Nakamura et al., 2016, Ma et al., 2019, and Yang et al., 2021^11,13,14^. The data from Nakamura et al., 2016, were reprocessed and remapped using STAR (v2.5.1b) with the reference genome ‘Macaca_fascicularis_5.0.96’ from Ensembl ^63^. Gene expression was quantified using RSEM (v1.3.0) with cell annotations downloaded from the original publications. Cell annotations for Yang et al., 2021, were downloaded from http://www.nhp-embryo.net, with raw reads kindly provided by the authors.

### Quality control and normalization

To filter out low-quality cells, we implemented a cut-off based on the number of expressed genes (nGene) and the percentage of mitochondrial genes (percent.mito). For Smart-Seq2 datasets, high-quality cells were required to have a minimum of 2000 genes expressed (nGene) and a percent.mito of less than 0.125. For 10X datasets or other datasets, the cut-off was determined from original publication. For the ones which still have too few nGene or too high percent.mito, the cutoff was further determined based on the general distribution of nGene and percent.mito. An upper limit for the number of nGene was set to prevent doublets in 10X datasets. Detailed parameters used in QC for each dataset are listed in the Table S4.

After quality control and the exclusion of mitochondrial genes, genes with expression in at least five cells were selected, assessed separately for each dataset. Subsequently, we calculated log-normalized counts using the deconvolution strategy implemented by the “computeSumFactors” function in the R scran package (v.1.14.6) ^64^, followed by rescaled normalization using the “multiBatchNorm” function in the R batchelor package (v.1.2.4) ^9^. This ensured that the size factors were comparable across batches. The log-normalized expression after rescaling was then utilized for human dataset integration, marker detection, and identification of differentially expressed genes.

### Restoration of previous cell annotations for published datasets

Original cell identities for human embryonic datasets, excluding those from Petropoulos et al, Xiang et al, and Tyser et al ^3–5^, were obtained from their respective original publications. We utilized the most recently published annotation from Meistermann et al ^7^ for the Petropoulos et al., dataset in our analysis. Using our embryonic reference, we observed misannotated cells from the Xiang et al., dataset, as also reported by Chhabra et al ^65^ (Figure S13A), and as such, used the annotations provided by Chhabra et al for Xiang et al. Further, the cross-species integration analysis in Yang et al ^11^ with the human CS7 data from Tyser et al ^5^, displayed that some of human advanced mesoderm cells overlapped with the cynomolgus monkey extraembryonic mesenchyme cells. In agreement with their observation, integration of the CS7 human gastrula together with postimplanted marmoset data ^15^, we observed that a proportion (53/159 cells) of the previously annotated human advanced mesoderm cells overlapped with the marmoset stalk cells. As such, 53 cells previously classified as advanced mesoderm cells were reannotated as “extra-embryonic mesoderm” cells, using the spatial data provided for “Stalk cells” from the postimplanted marmoset ^15^ (Figure S13B) during cross-species integration (created by the “RunFastMNN” function with 2000 top variable genes and 25 top PCs). We then performed a DEG analysis between the remaining advanced mesoderm cells and extra-embryonic mesoderm cells and observed that the 53 cells displayed intermediate expression for both these lineages, providing additional evidence supporting the reannotation (Figure S13C). Additionally, a total of 330 EVT and 649 STB cells were identified from previously annotated CTB cells in the cynomolgus monkey (Macaca fascicularis) ^11,13,14^, and 51 CTB and 77 STB cells in the Marmoset ^15^ and confirmed by the expression of markers from human CTB, STB, and EVTs (Figure S13D-G). TrB cells from Ai et al ^29^ were selected using the Seurat pipeline (with 2000 top variable genes and 25 top PCs) to further delineate the CTB, STB, and EVT lineages. For datasets lacking cell annotations, we attempted to restore annotations based on their source code or descriptions provided in the methods section of original publication. All reanalyzed cell annotations were further validated by the marker gene expression reported in the original publication.

### Construction of the human embryonic reference

The human embryonic reference was established by integrating previously published datasets, which included six sets of human embryonic data spanning zygote early embryos, *in vitro* cultured human blastocysts, 3D *in vitro* cultured human blastocysts up to pre-gastrulation stages, and a Carnegie CS7 (16-19 dpf) human gastrula. The integration process utilized the fastMNN calculation method from the batchelor (v1.6.2) package. In detail, the top 4000 variable genes were initially selected using the “SelectIntegrationFeatures” function from Seurat package. It is noteworthy that four out of the six reference datasets were preimplantation datasets. To mitigate voting bias during the “SelectIntegrationFeatures” function, the top 2000 variable genes were first selected from the four preimplantation datasets. Subsequently, a combination consideration was applied to the remaining post-implantation dataset and CS7 dataset. The linear merge order of batches was determined by the order of embryo developmental timepoints represented in the datasets. After obtaining the MNN-corrected PCA subspace results from the fastMNN calculation, the Uniform Manifold Approximation and Projection (UMAP) dimensional reduction result was calculated using the “umap” function from the uwot package (v0.1.14) (https://CRAN.R-project.org/package=uwot), employing the top 50 MNN-corrected PCA subspace. The entire dataset was clustered using the Leiden algorithm (“RunLeiden” function from the Seurat package) ^66^, utilising the same neighbourhood graph constructed from the corrected PCA subspace. Epiblasts belonging to clusters C1 and C10, 12 were identified as early and late epiblasts, respectively. Primitive endoderm (PE) cells were split as early (clusters C1 and C22) and late hypoblasts (cluster C11), respectively (Figure S1B and C). Throughout this process, the grand mean values (grand.centers) values, singular value decomposition results (unitary matrix) from MNN corrections, variation along the batch vectors, UMAP model, and cell clustering were recorded for query dataset projection and identity prediction.

### Identification of marker genes and regulatory activity of transcription factors within human embryonic datasets

Lineage marker genes were identified using the “FindAllMarkers” function with default parameters, setting the adjusted p-value cutoff at less than 0.05. To identify the markers conserved in primate, marker detection was performed within each species and p-values were combined using “stouffer” function from poolr package(v1.1-1) ^67^ and followed by adjustment. The categories hypoblast, definitive endoderm, and yolk sac endoderm are collectively referred to as “Endoderm,” while preimplantation TE (Trophectoderm), CTB (Cytotrophoblast), STB (Syncytiotrophoblast), and EVT (Extravillous trophoblast) are grouped as “TEs”. Additionally, “ExE_Mech,” “Stalk,” and “ExE_Mes” are consolidated as the “ExE_Mes_stalk” group, representing the extraembryonic lineage. AdvMes (Advanced Mesoderm) and axial mesosderm from humans have been excluded from the mesoderm groups due to their similarity with extraembryonic lineages or distinction from the main mesoderm. “EmDisc” from marmoset has been included in the epiblast group as the majority of them overlap with the epiblast. The pseudotime trajectory was computed using the R package slingshot (v.2.6.0) ^68^. This package facilitates the computation of lineage structures in a low-dimensional space. In summary, pre-computed cell embeddings and annotations obtained from a human embryonic reference served as input for the “slingshot” function. The start cluster was set to “zygote,” followed by the application of the “slingPseudotime” function to infer individual pseudo time. The cells, categorized into prelineages (zygote, 2-4 cell, 8 cell, morula, and E5 prelineage), ICM, epiblast, hypoblast, TE, and CTB, were utilised to infer the main trajectories. The DynamicHeatmap function from the R package SCP (v0.5.6, available at https://github.com/zhanghao-njmu/SCP) was employed to identify transcriptional factor genes significantly associated with the pseudotime in the three main trajectories start from themorula stage. For a more detailed Epiblast-PriS-Mesoderm-Amnion trajectory, cells belonging to prelineages, epiblast, PriS, mesoderm, amnion, and AdvMes cells were included. Similarly, cells belonging to prelineages and hypoblast were considered to infer the detailed hypoblast trajectory. In the case of the TE-CTB-EVT-STB trajectory, cells belonging to prelineages, TE, CTB, STB, and EVTs were included. Batch-corrected gene expression from six human embryonic datasets was initially calculated using the “mnnCorrect” function from the batchelor package. The resulting expression profiles were then utilized to identify regulatory modules by inferring co-expression with transcription factors through the “pyscenic grn --method grnboost2” command. Each coexpression module served as input for cis-regulatory motif analyses conducted by running “pyscenic ctx” with the following motif collections: “hg38 refseq-r80 10kb_up_and_down_tss.mc9nr.feather,” “hg38 refseq-r80 500bp_up_and_100bp_down_tss.mc9nr.feather,” and “motifs-v9-nr.hgnc-m0.001-o0.0.mod.tbl”^22^. Subsequently, the AUC (area under the curve) values for each regulon were computed using “pyscenic aucell”. Valid regulons were further refined by applying filters, requiring regulon activity in more than 1% of total input cells and an average AUC value exceeding 0.1 for cells belonging to the same lineages.

### Cross-species integration

The integration of human embryonic datasets with scRNA-sequencing data from cynomolgus monkeys and marmosets included following steps. First, cynomolgus monkey gene IDs were converted to Homo sapiens gene symbols using the ortholog list from Ensembl ^69^. Marmoset genes were merged based on shared gene names within human dataset. Subsequently, rescaled normalization using the “multiBatchNorm” function was performed separately for each species. Cells from Yang et al were down sampled to 2000 ^11^. The top 3500 variable genes for cynomolgus monkeys and marmosets were selected using “SelectIntegrationFeatures” function, and cross-checked with the top variable genes used in the human embryo reference to confirm overlapping genes amongst all three species construction. Among the 14,978 genes shared by all three species, a total of 6005 top variable genes were identified. Of these, 544 or 1597 genes were identified as the top variable genes in all three species or in any two species, respectively. Additionally, the top 500 species-unique variable genes were included, resulting in a final selection of 3641 genes for the integration of all three species. To preserve the specificity of each individual species while emphasizing commonalities, the “mnnCorrect” function from the batchelor package was used to calculate batch-corrected expression within each species separately. The batch-corrected expression of the three species, which removed within-species batch differences, served as input for the “FindIntegrationAnchors” and “IntegrateData” functions from the Seurat package for integration. Subsequently, the top 20 principal components were calculated using the “RunPCA” function and utilized for UMAP dimensional reduction with the “RunUMAP” function.

For the integration of scRNA-sequencing data from the preimplantation blastoids with that from *in vitro* cultured post-implantation cynomolgus monkey embryos (wild-type, Day14) ^11^, normalization was performed separately for each dataset. Each human blastoid dataset and cynomolgus monkey dataset were then integrated using the “RunfastMNN” function (wrapped in the R SeuratWrappers package) with 2000 anchor features and the top 25 principal components. The resulting components were used for UMAP dimensional reduction with the “RunUMAP” function.

### Projection of query dataset on human embryonic reference

To project the query dataset without influencing original dimensional reduction of reference dataset, the following steps are taken:

1. Normalization: when incorporating a query dataset, it is rescaled to the lowest-coverage batch of the reference dataset ^6^. The rescaling factor used for the reference dataset is also applied to the query dataset.
2. MNN Correction and PCA projection: implementing MNN correction to remove batch effect assumes the presence of mutual nearest neighbors (MNNs) define the most similar cells of the same type across batches and the batch effect is almost orthogonal to the biological subspace. In our reference dataset, distinguishing between two cell populations,TE and amnion, poses challenges due to their gene expression similarities. Additionally, these populations exhibit an extremely uneven distribution (almost 39:1) across our six datasets, with only around 30 amnion cells included in the reference dataset. To address potential errors in MNN pairs, especially in datasets with a high proportion of amnion/aminon like cells or TE/TE like cells, we divided the query dataset into several maximal 200 subsamples to avoid MNN pair saturation. Each sample was randomly selected five times to form different subsamples, preventing bias introduced by downsampling. MNN calling was performed using the “findMutualNN” function between the query dataset and each dataset separately, utilizing cosine normalization values. The default value of k was set to 20, except when the number of query samples was less than 50, where k was set to 5. To further enhance accuracy, query cells assigned to have both amnion and TE reference MNN pairs were discarded in downstream analysis. Subsequently, the query cosine-normalized data underwent removal of the same grand mean values (grand.centers) of each gene from reference dataset construction. This was followed by a dot product calculation with left singular vectors(unitary matrix) from singular value decomposition in reference construction and removing variation along the reference batch correction vector(Orthogonalization) using the internal functions “.orthogonalize_other” and “.center_along_batch_vector”, previously wrapped in the fastMNN() function, placed the query dataset into the same corrected batch-corrected PCA space as the reference datasets. As described in the original paper ^9^, cell-specific batch correction vectors were calculated based on identified MNN pairs, and a further batch correction on PCA space was performed using internal functions “.compute_correction_vectors” and “.adjust_shift_variance”, previously wrapped in the “mnnCorrect” function from batchelor(v1.6.2) ^9^.
3. UMAP Calculation: after obtaining batch-corrected PCA subspace results for the query dataset, two UMAP projections were calculated using the “umap_transform” function based on the previously mentioned reference UMAP model.

To filter out non-related cells, a Spearman correlation was calculated between the query data and the reference dataset using cosine normalization values. To set the threshold for filtering irrelevant cells, processed expression files from unrelated datasets, including five scRNA-seq datasets from human pancreas, macrophage, fetal kidney development, and liver tissues, were downloaded from https://cblast.gao-lab.org/ to test the correlation coefficient with human embryonic reference cells ^70–74^. In our analysis, cells with a median top correlation coefficient (calculated within top 20 correlated reference cells) less than 0.5 were considered non-related. In addition, before the entire projection calculation, raw counts for cells from 10X datasets stratified by different timepoint with similar expression patterns were aggregated within neighborhood nodes, as calculated by miloR package(v1.2.0) ^75^. This aggregation enhances our MNN calculation and correlation calculation.

### Cell prediction based on UMAP projection

To predict cell identities based on UMAP projection, we initially constructed lineage-representing polygon regions on the reference UMAP using the “chull” function from the ‘grDevices’ R package(v4.0.3) (Figure S1B) (https://www.R-project.org/). In this process, reference cells that did not fall into the main corresponding lineage clusters were removed. The epiblast and hypoblast polygon regions were further split into early/late epiblast and hypoblast regions based on cluster information, confirmed by the timepoint distribution and DEG expression (Figure S1D).

Query cells (or neighborhood from miloR calculation) that passed the correlation filtering and were projected onto the reference UMAP were first checked for intersection relationships with reference polygon regions using the ‘st_intersects’ function from the ‘sf’ (v.0.9-7) package^76^. For those overlapping with multiple reference polygon regions (e.g., overlapped polygon regions belonging to advanced mesoderm cells with extraembryonic mesoderm cells), all reference lineages would be reported. For those not falling into constructed polygon regions, a k-NN approach was utilized to annotate query cells with reference data by calculating the Euclidean distance in the UMAP. For each query cell, the 5 nearest neighbors from the reference cells were selected, and the cell type or developmental time was predicted by majority vote. During this process, based on the assumption that MNN pairs represent the most similar cells of the same type across batches, predicted lineages without support from MNN pairs were labeled as “ambiguous” possibly due to inaccurate UMAP transformation or projection for query cells. For query datasets aggregated with neighborhood nodes, cells contributing to the neighborhood were assigned the same predicted annotation as that for the neighborhood and multiple predicted lineages were reported by majority vote also. Cells that failed to form a neighborhood during miloR calculation were labeled as “nb_failed”.

### Module score calculation

The predicted lineage scores for each cell were calculated using the “AddModuleScore” function from Seurat package, incorporating the top 15 identified lineage marker genes.

### Detection of top DEGs between amnion, primitive streak and extraembryonic mesoderm cells and TE

Gene expression of amnion, primitive streak, and extraembryonic cells, from Tyser et al ^5^ were compared with pre- and post-implantation TE cells from Petropoulos et al., Yanagida et al., Meistermann et al., and Xiang et al ^3,4,6,7^. The TE cells from Xiang et al. were categorized into pre-implantation TE and post-implantation CTB, STB, and EVT. “FindMarkers” function from the Seurat package was employed for differential expression analysis, utilizing the ‘wilcox’ test to compare all TE populations with amnion, primitive streak, or extraembryonic mesoderm cells. Since cells of late lineages and TE lineages were not from the same dataset, batch difference could influence DEG detection. For a gene to be considered truly differentially expressed, it had to meet four additional criteria: (1) Average expression level in up-regulated lineages greater than 10. (2) Log2(fold change) greater than 0.25 in all five comparisons (compared with all five TE groups). (3) FDR(False Discovery Rate) less than 0.05 in at least four comparisons. (4) Additionally, each gene had to meet the following conditions in at least five comparisons: Percentage of cells expressing the gene in the highly expressed group greater than 50%. Percentage of cells expressing the gene in the lowly expressed group less than 25%. To ensure consistency, DEGs located on the sex chromosome were excluded from analysis, considering only the male embryo included in Tyser et al ^5^.

### Detection of DEGs between Naïve/ Primed derived TLC and preimplantation embryonic TE

The gene expression profiles of predicted naïve and primed TE-like cells were compared with TE cells obtained from studies by Petropoulos et al., Yanagida et al., Meistermann et al., and Xiang et al. ^3,4,6,7^, specifically focusing on pre-implantation TE cells (Day 6 to Day 7). Differential expression analysis was conducted using the “FindMarkers” function from the Seurat package and the ‘wilcox’ test. When comparing all TE populations, considering that the non-embryonic datasets were all generated on 10X platforms, for a gene to be considered truly differentially expressed, it needed to meet three additional criteria: (1) Log2(fold change) > 0.25 in at least four comparisons. (2) FDR < 0.05 in at least three comparisons. (3) Percentage of cells expressing the gene in the highly expressed group > 50%, and expressed percentage in the lowly expressed group < 25% in at least three comparisons.

### Detection of DEGs between blastoids derived ELC, HLC TLC vs preimplantation embryonic correlative lineages

Gene expression profiles of ELC, HLC, and TLC from studies by Yanagida et al., Kagawa et al., and Yu et al ^6,32,45^ were compared with preimplantation embryonic correlative lineage cells from studies including Petropoulos et al., Yanagida et al., Meistermann et al., and Xiang et al., respectively ^3,4,6,7^. Differential expression analysis was conducted using the “FindMarkers” function and the ‘wilcox’ test, comparing blastoid lineage-like cells with embryonic reference cells. Given the limited number of reference cells for the preimplantation hypoblast in studies by Yanagida et al., Meistermann et al., and Xiang et al. (14, 2, and 7 cells, respectively) ^4,6,7^, hypoblast cells from Meistermann et al were excluded in the reference comparison. Comparisons between HLC and PE cells from Yanagida et al. and Xiang et al. were performed using the ‘roc’ test, with a power greater than 0.8 considered significant. To be considered truly differentially expressed, a gene had to meet two additional criteria, including being a log2(fold change) > 0.25 in all four comparisons, having an FDR < 0.05 in at least three comparisons.

### Web Interface

A shiny app generated by shinyCell ^59^, which includes the above integration, as well the web interface for prediction can be browsed at http://petropoulos-lanner-labs.clintec.ki.se.

## Data and code availability

In-house primed-derived TLC ScRNA-seq data have been deposited at GEO and are publicly available as of the date of publication. Accession numbers are listed in the Table S4. All original code has been deposited at GitHub is publicly available as of the date of publication. Any additional information required to reanalyze the data reported in this work paper is available from the lead contact upon request.

## Table legend

Table S1: Transcription factors associated with epiblast, hypoblast and TE trajectories. Table S2: Lineage markers genes identified in human early embryos.

Table S3: Conserved lineage markers genes identified in human, cynomolgus monkey and marmoset.

Table S4: Datasets used in this analysis.

Table S5: DEGs identified between naïve and primed cells and between preimplantation and late lineages of embryonic TE cells.

Table S6: DEGs identified comparing naïve and primed-derived TLC with and preimplantation embryonic TE cells.

Table S7: DEGs identified between early and late epiblast, early and late hypoblast, and amongst TE, CTB, EVT and STB lineages.

## Acknowledgements

We thank members of the laboratories of F.L. and S.P. for discussions. This work was supported by the Swedish Research Council (F.L., S.P.), Ragnar Söderberg Foundation (F.L.), Ming Wai Lau Center for Reparative Medicine (F.L.), Center for Innovative Medicine (F.L.), Wallenberg Academy Fellow (F.L.), Swedish Society for Medical Research (S.P.), Emil och Wera Cornells Stiftelse (S.P.), Natural Sciences and Engineering Research Council of Canada Discovery Grant (S.P), The Canadian Institutes of Health Research (S.P., J.R.), Päivikki and Sakari Sohlberg Foundation (J.W.), and Sigrid Jusélius Foundation (J.W.). SP holds the Canada Research Chair in Functional Genomics in Reproduction and Development.

We would like to thank Dr Elmir Mahammadov, Dr Rang Yang and Dr Alexander Goedel, Professor Shankar Srinivas, Professor Matteo Mole, Professor Magdalena Zernicka-Goetz, Professor Antonio Scialdone, Professor Yu Yi, Professor Zongyong Ai, Professor Tianqing Li for sharing with us their processed gene expression matrices and corresponding published annotation from their work.

The computations were partially enabled by resources in project SNIC 2022/22-786, SNIC 2022/22-786 and SNIC 2020/15-102 provided by Uppsala University at UPPMAX.

## Contributions

C.Z., S.P., and F.L. conceived the study with advice from A.P-R., and J.P.S.. A.P-R., J.W., produced primed derived TLC which were sequenced with assistance from L.B.V.. Data analysis was performed by C.Z. with input from S.P., F.L., and Å.B. Interpretation of results was performed by C.Z., F.L., S.P., A.P-R., J.P.S., J.W., N.M.O. Y.Z., B.C., J.R., and J.F. The manuscript was written by C.Z., F.L., and S.P., with input from all of the authors.

## Ethics declarations

### Competing interests

The authors declare no competing interests.

---

**Figure S1.**
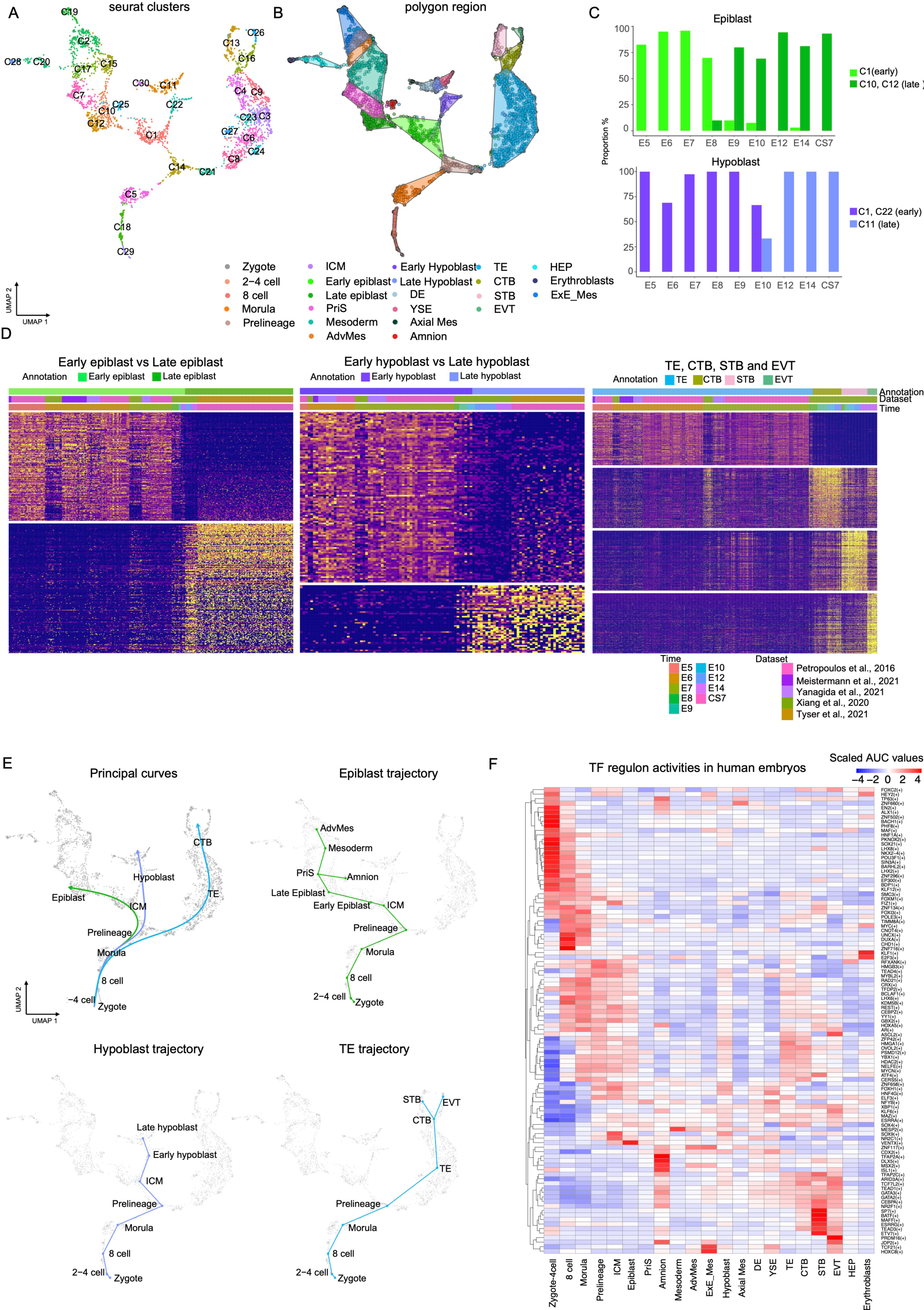
Clusters, trajectories and regulon activity within the human embryonic reference. (A) Unassigned cluster distribution of UMAP used in Figure 1A. (B) Polygon regions constructed based on lineage annotation and clusters. (C) Cell distribution in clusters for epiblast and hypoblast cells. (D) Heatmap showing expression of DEGs between early and late epiblast, early and late hypoblast and DEGs among TE, CTB, STB and EVT. (E) Principle curves, and trajectories constructed from slingshot. (F) Heatmap displaying average AUC values of enriched regulons within each lineage.

**Figure S2.**
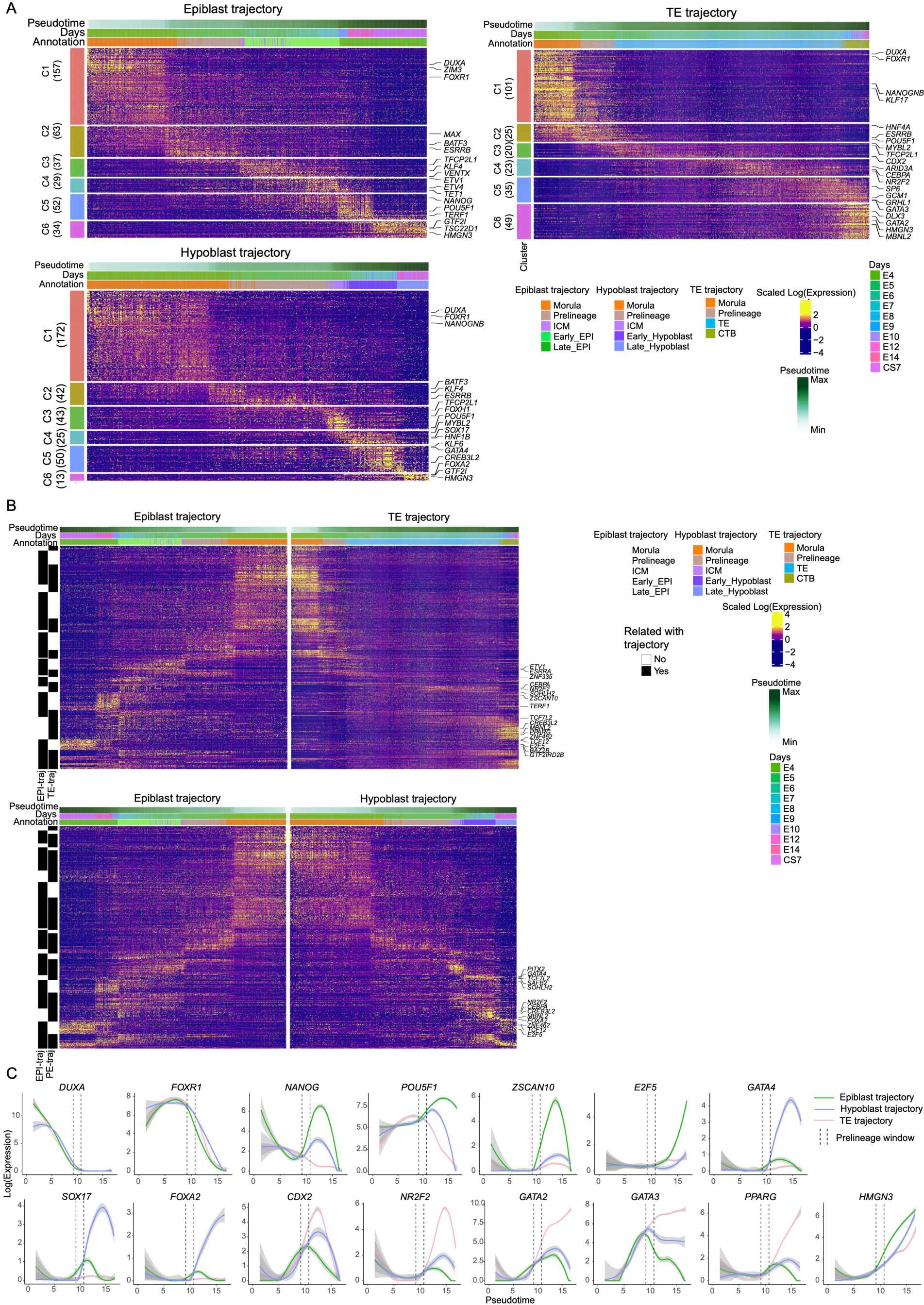
Transcription factor gene expression along the epiblast, hypoblast, and TE trajectories. (A) Heatmap of expression of transcription factor (TF) genes which were significantly related to trajectories pseudotime. Cluster pattern of expression was indicated on the left with numbers indicating the number of TF genes. (B) Joint heatmap showing expression of TF genes related to epiblast/TE trajectories and epiblast/hypoblast trajectories. The black and white annotation on the left indicated whether corresponding TF were significantly related to pseudotime. (C) Expression dynamics (pseudotime) of selected transcriptional factor genes along 3 main trajectories. The confidence interval (95%) is indicated by bandwidth. Different trajectories are indicated by colours, respectively.

**Figure S3.**
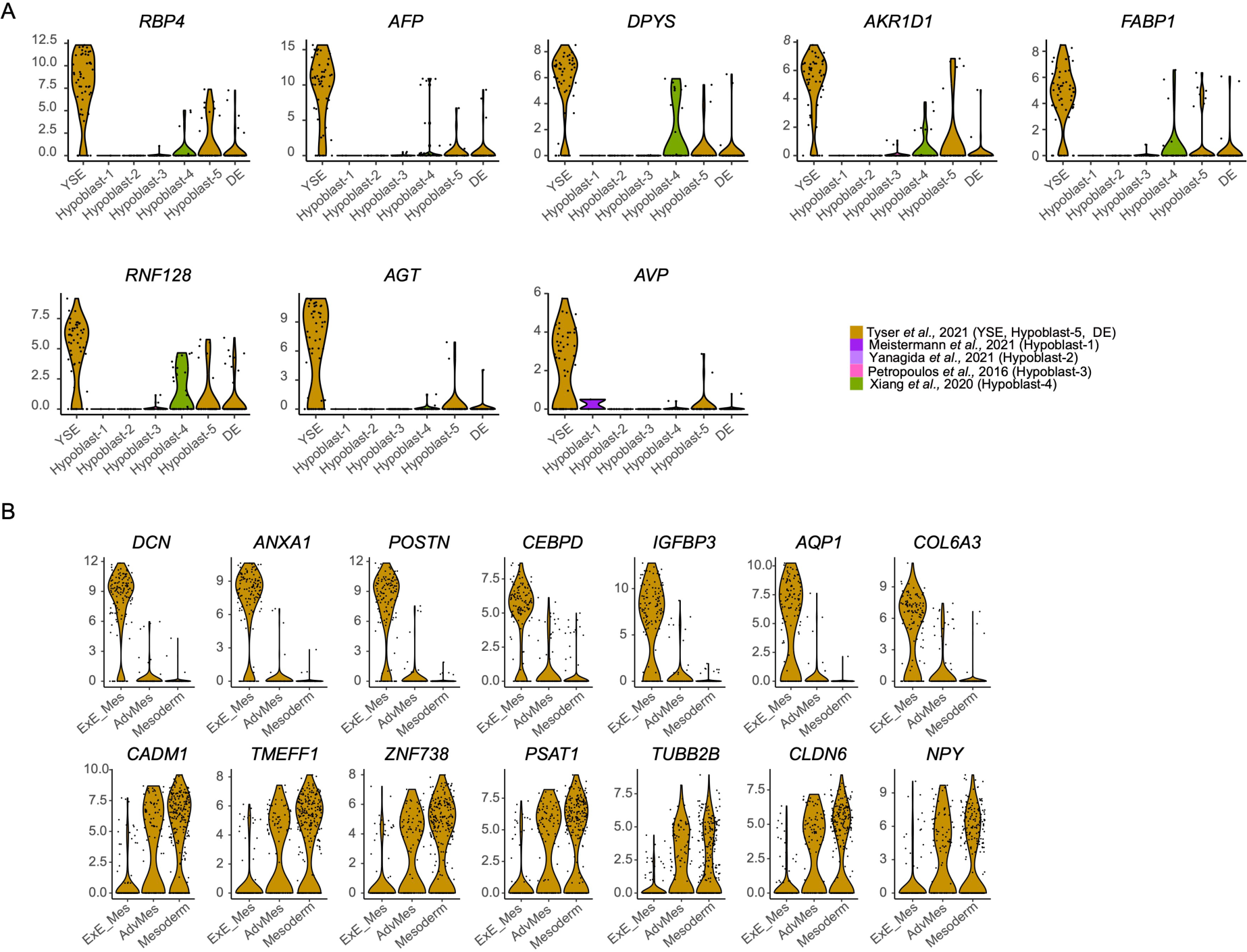
Violin plot showing distinct markers expressed in YSE in comparison to hypoblast and DE (A), and top DEGs between ExE_Mes and embryonic mesoderm cells (B).

**Figure S4.**
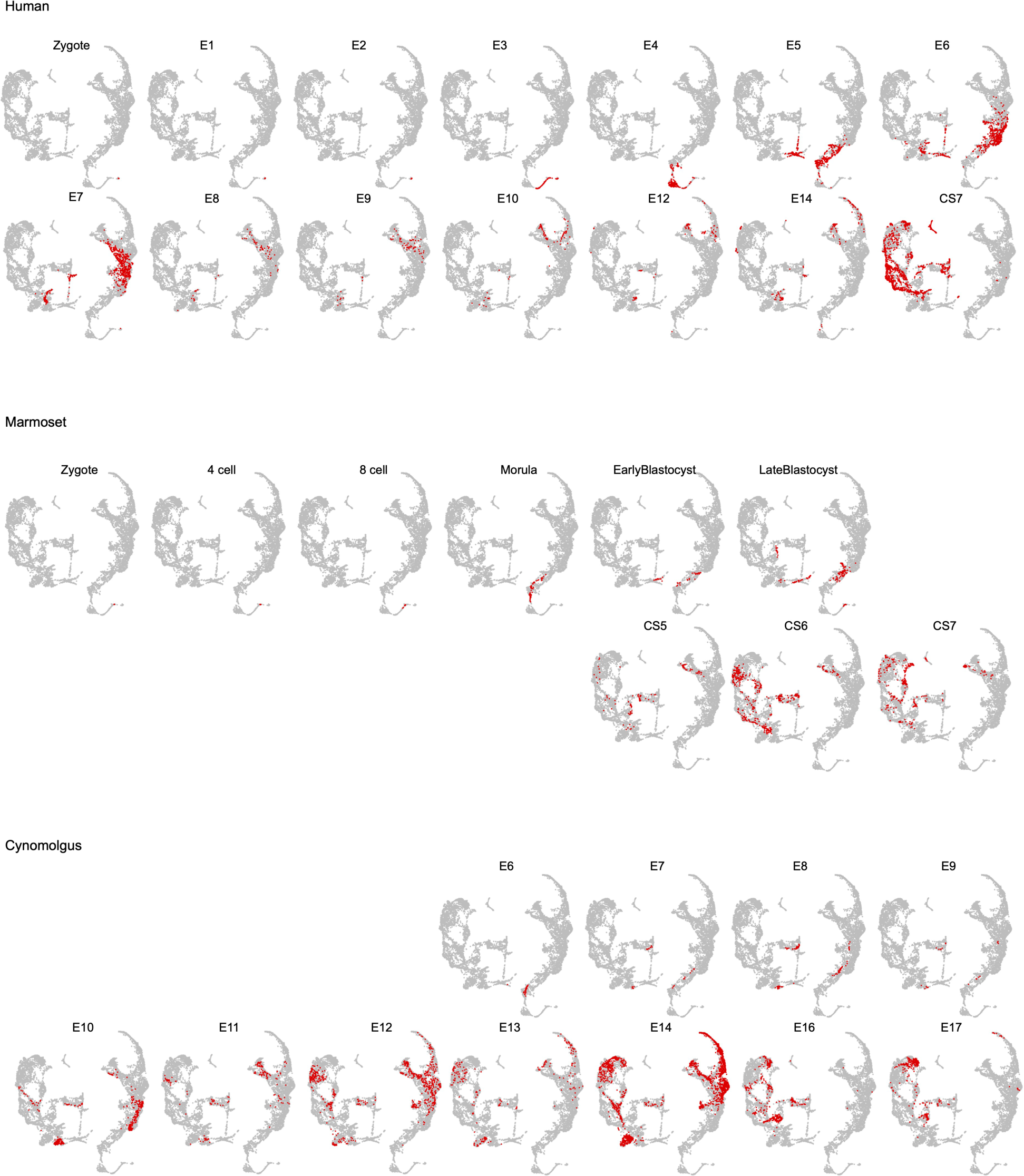
Primate cross-species integration of embryonic datasets with cells highlighted from the different embryonic time points for each species.

**Figure S5.**
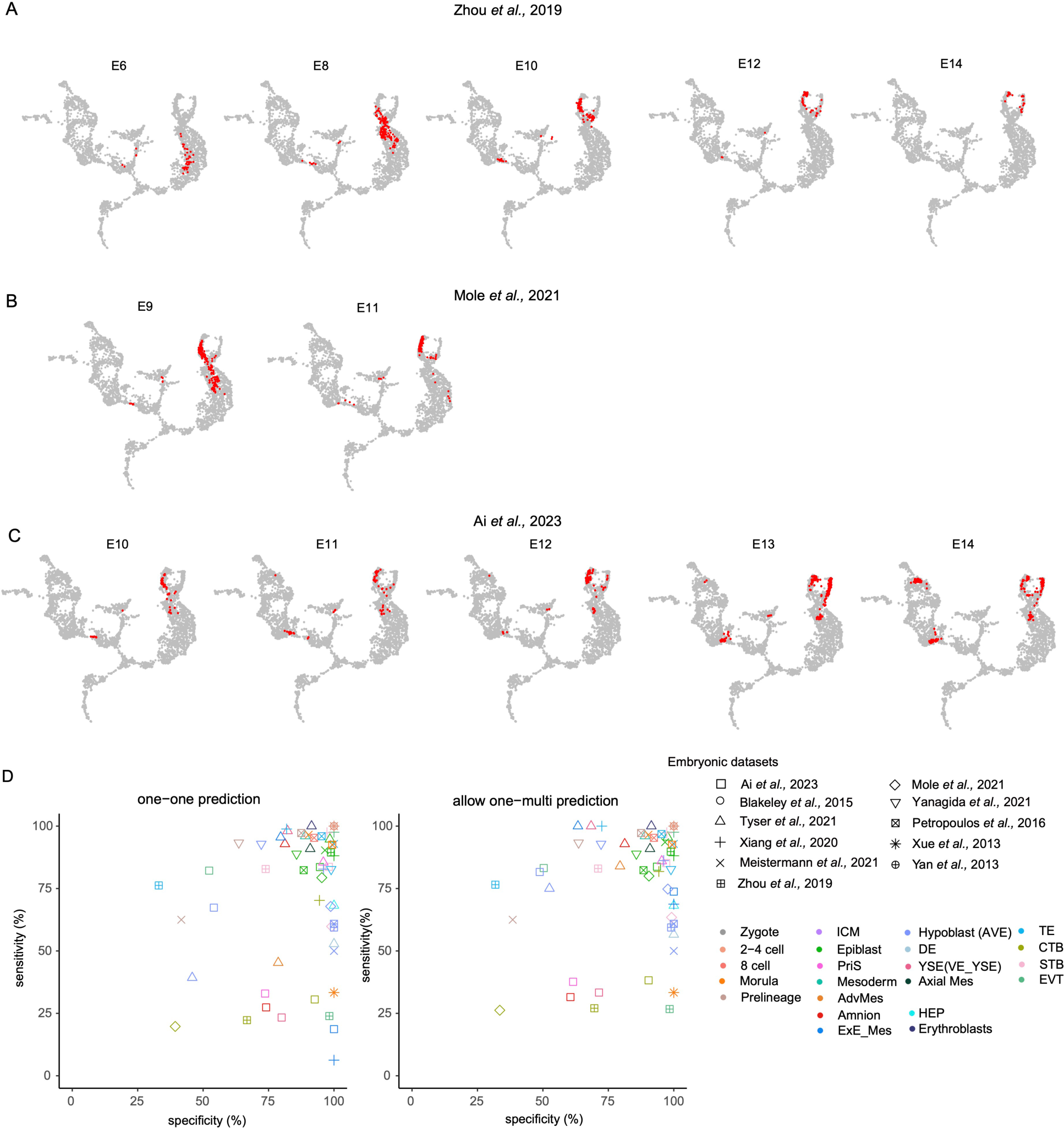
(A, B, C) Highlighted cells (neighbourhood nodes) from different embryonic time points for zhou et al (A), Mole et al (B) and Ai et al (C) from the projection on embryonic reference. (D) Evaluation of predicted sensitivity and specificity for each cell type in the embryonic datasets. The shape and colour of data points indicate queried cell types and data sources, respectively. Left) Allowing one-to-one predictions; a single prediction with multiple reference lineages was treated as misannotated and not true. For example, if “CTB” cells were only predicted as “CTB”, this was considered a true prediction. Right) Allowing one-to-many predictions; which consisted of a single prediction with multiple reference lineages were treated as valid predictions. For example, if “CTB” was predicted as “CTB” or “CTB” plus any other lineage, it was considered to be a true prediction.

**Figure S6.**
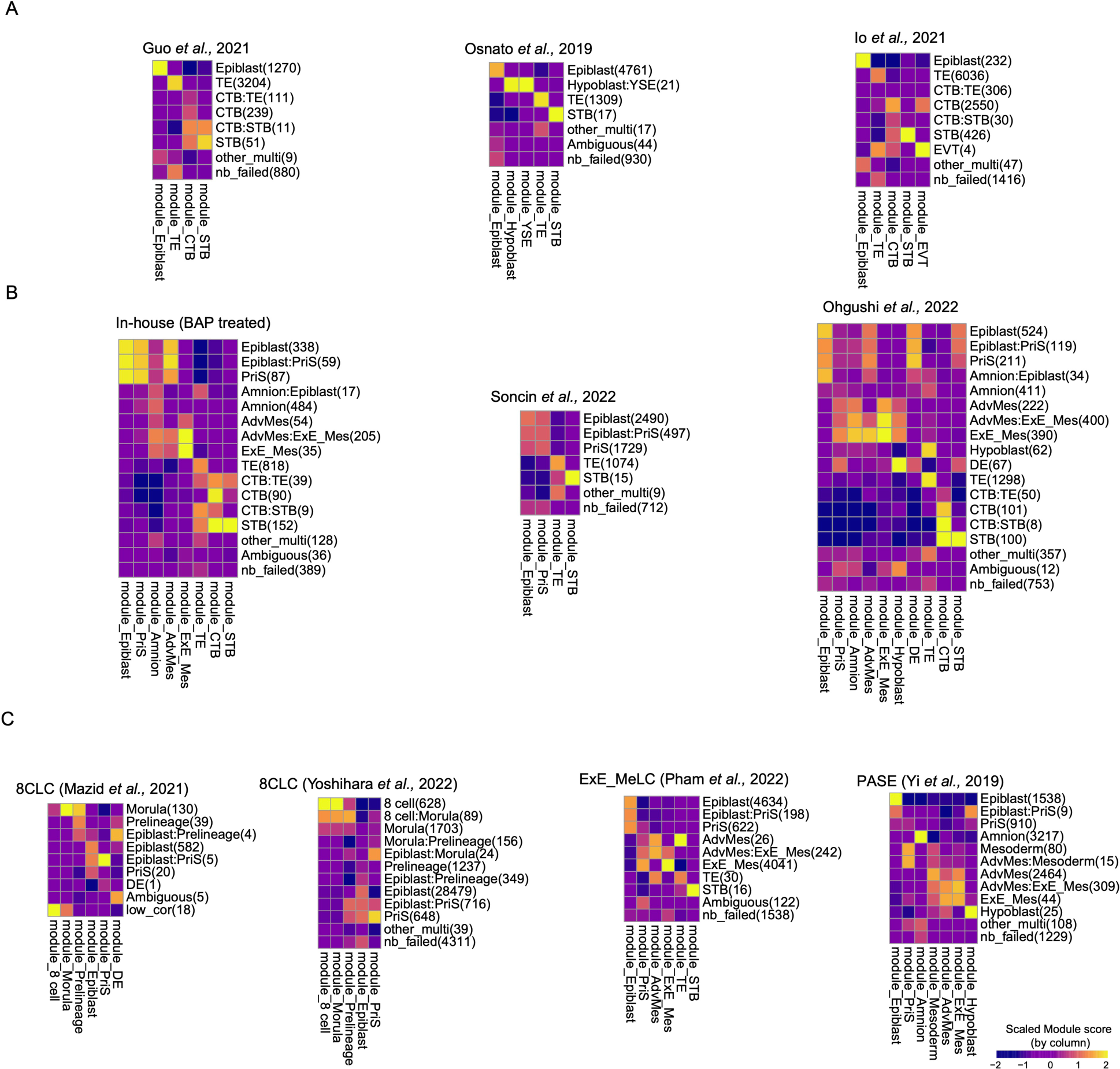
Module scores of corresponding predicted lineages in three naïve stem cells models (A), three primed stem cells models (B), two 8-cell-like models, one extraembryonic mesoderm-cell-like model, and PASE models (C). Columns represent different lineage models scores, and rows represent predicted lineages for each dataset. The predicted cell numbers are included in parentheses.

**Figure S7.**
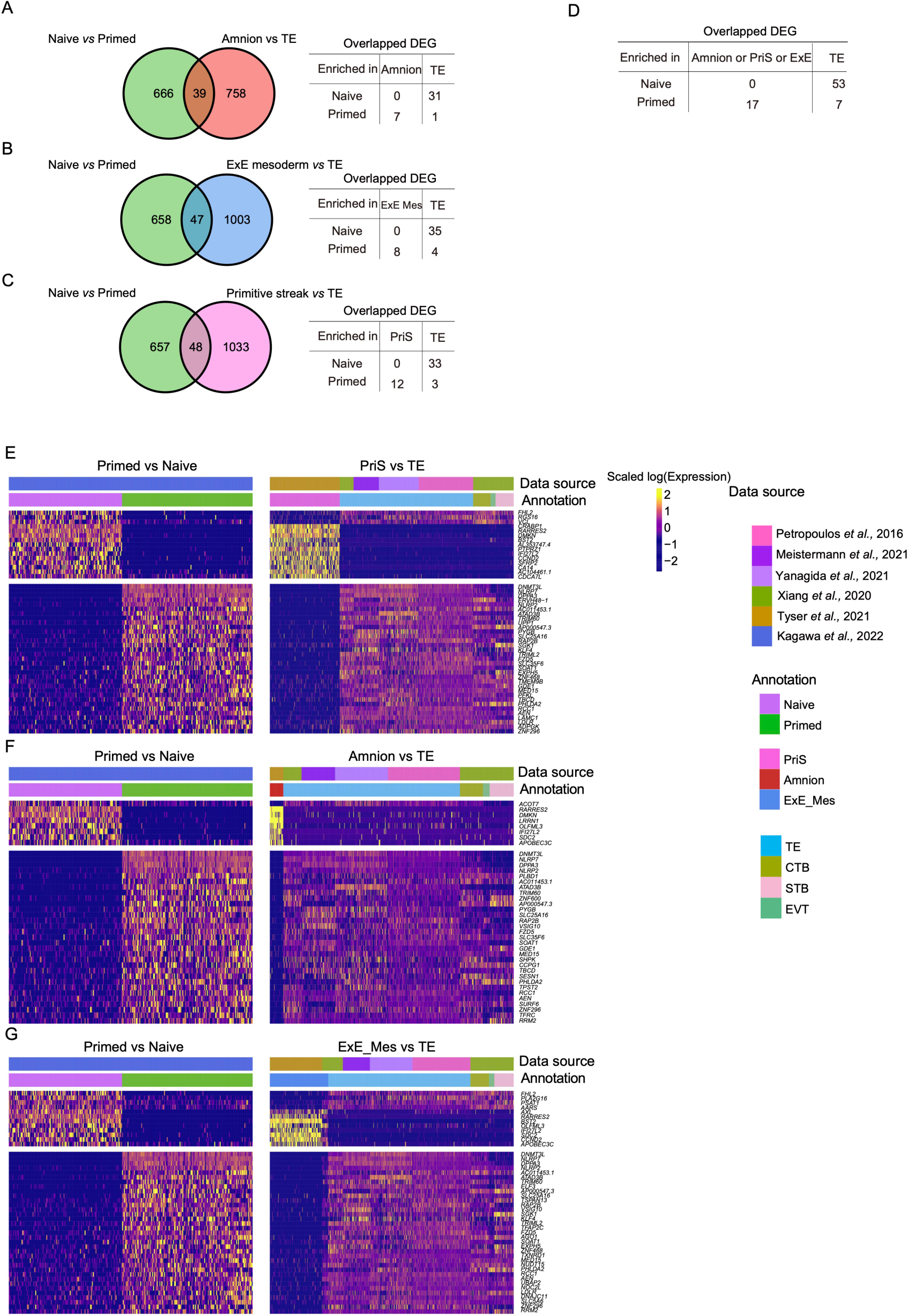
(A) Venn diagram showing overlaps between naïve vs. primed DEGs and amnion vs. TE DEGs. The detailed number of DEGs with direction is included in the tables on the right. (B) ExE_Mes vs. TE DEGs, and (C) PriS vs. TE DEGs. (D) The number of overlapping DEGs between late lineages vs. TE and naïve vs. primed. (E) Heatmap showing expression of DEGs identified in naïve vs. primed and PriS vs. TE, (F) DEGs shared by naïve vs. primed and amnion vs. TE, and (G) DEGs shared by naïve vs. primed and ExE_Mes vs. TE.

**Figure S8.**
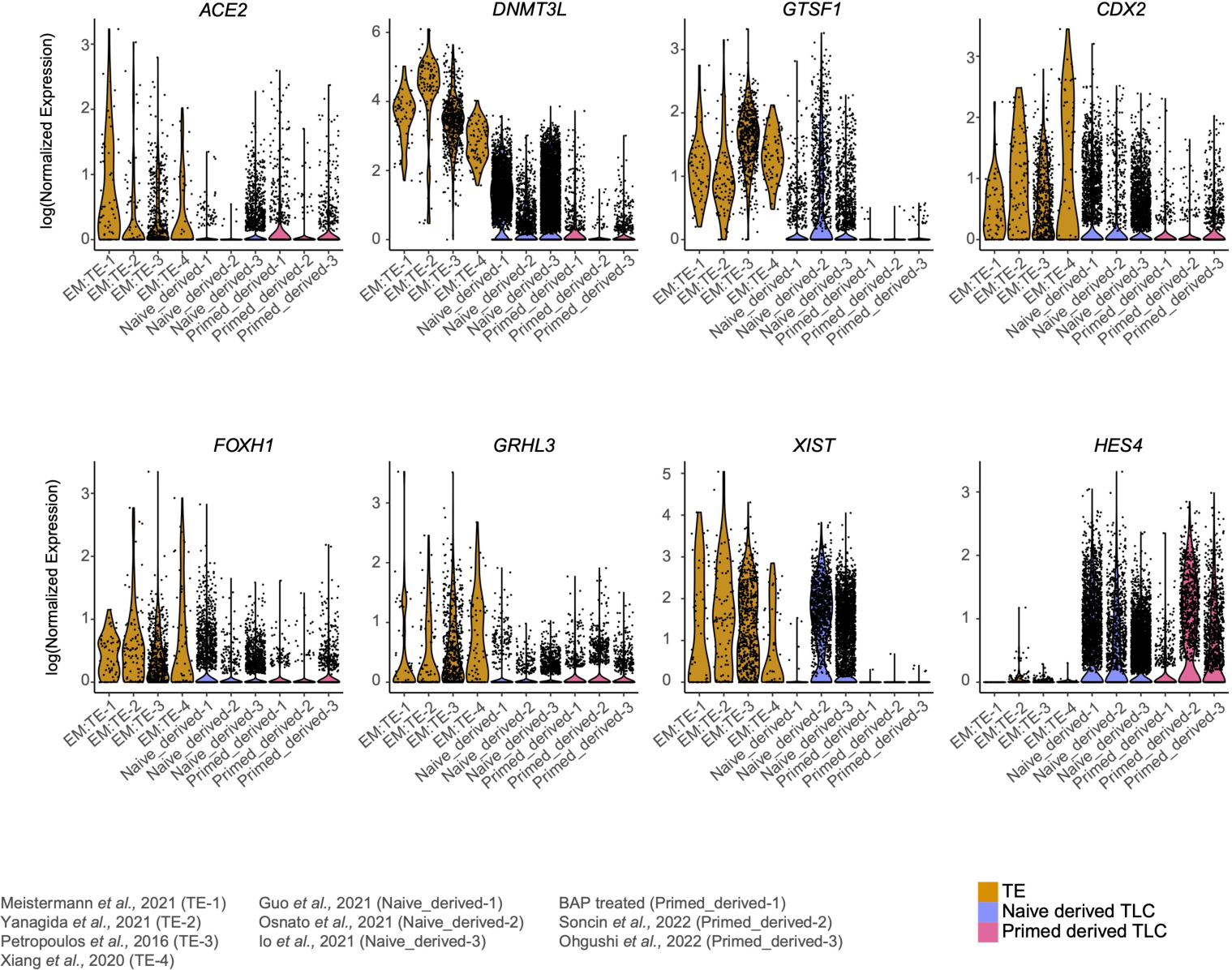
Violin plot showing representative DEGs between naïve or primed derived TLCs and embryonic preimplantation TE cells.

**Figure S9.**
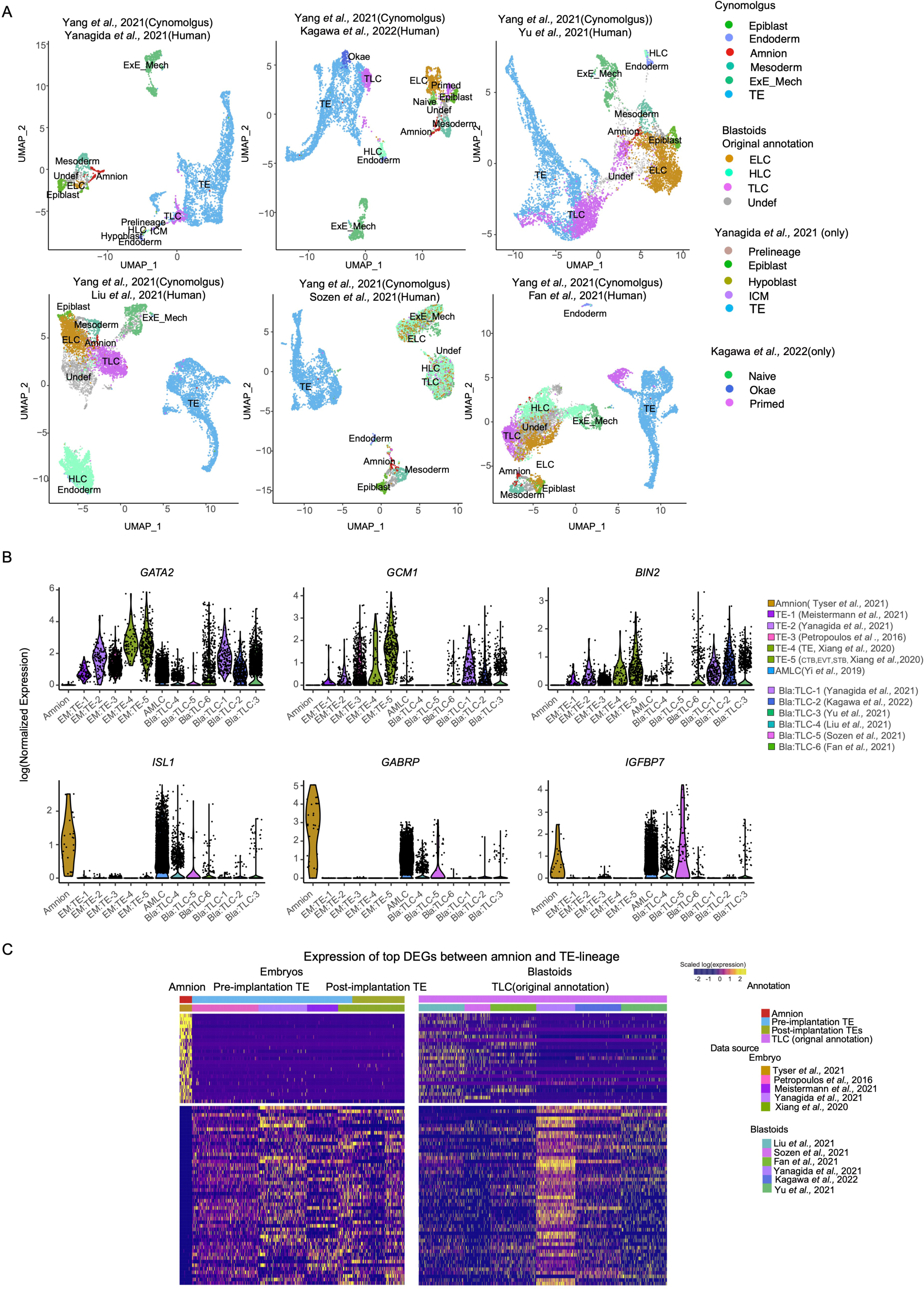
(A) UMAP projection of Mutual Nearest Neighbours (MNN) cross-species integration, including cells from each blastoid dataset and cynomolgus macaque embryonic dataset at Day 14 (Yang et al., 2021), coloured by cell type. Embryonic cells from Yanagida et al. and naïve, primed, and Okae cells from Kagawa et al. were also included in the integration. (B) Violin plots showing log-transformed expression of key amnion and TE markers in amnion, TE, amnion-like cells (AMLC), and TLC from the six blastoids (based on their original annotation). (C) Expression of significantly differentially expressed genes between amnion and TE in embryonic amnion, TE, and TLC from the six blastoid models. For visualisation, cells from the same cell type were randomly down-sampled to 200.

**Figure S10.**
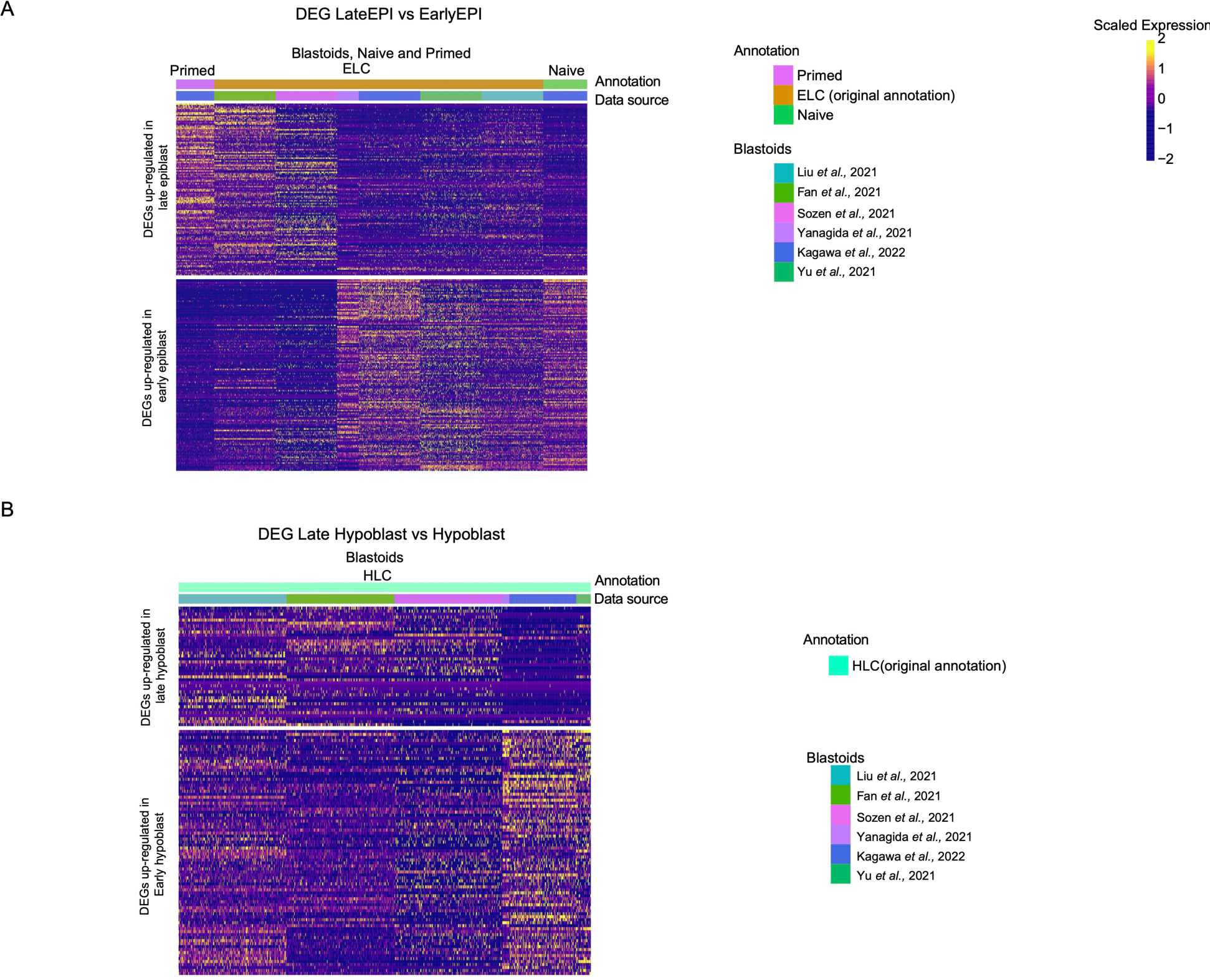
(A) Heatmap displaying DEGs between early and late epiblast in primed cells, blastoids ELC cells (based on their original annotation), and naïve cells. (B) Heatmap displaying DEGs between early and late hypoblast blastoids HLC cells (based on their original annotation).

**Figure S11.**
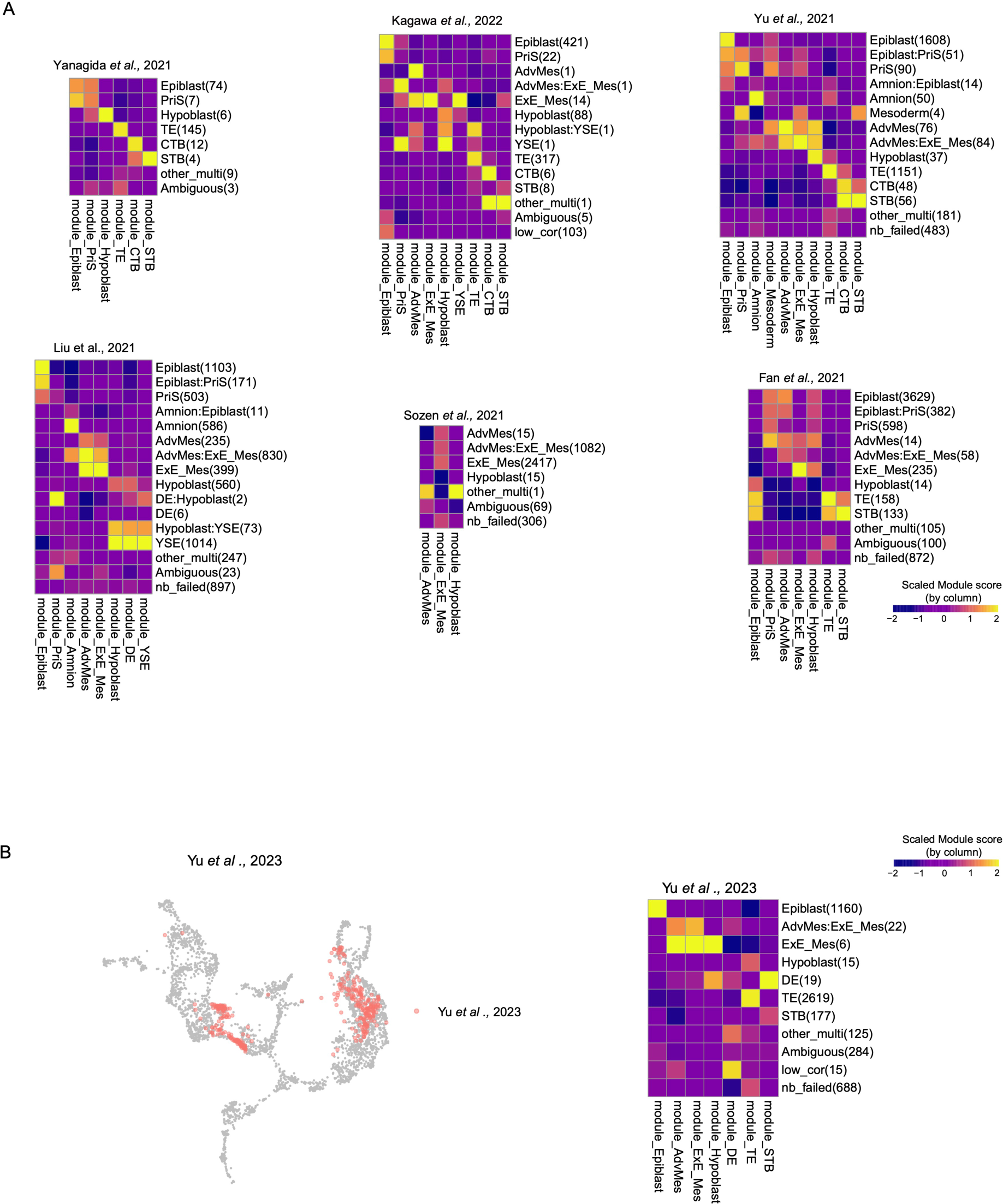
(A) Module score of corresponding predicted lineages in six blastoids. Columns represent different lineage models scores, and rows represent predicted lineages for each dataset. The predicted cell numbers are included in parentheses. (B) Projection of most-recent blastoids from Yu et al., 2023 onto human embryonic reference. Module score validation of predicted lineages.

**Figure S12.**
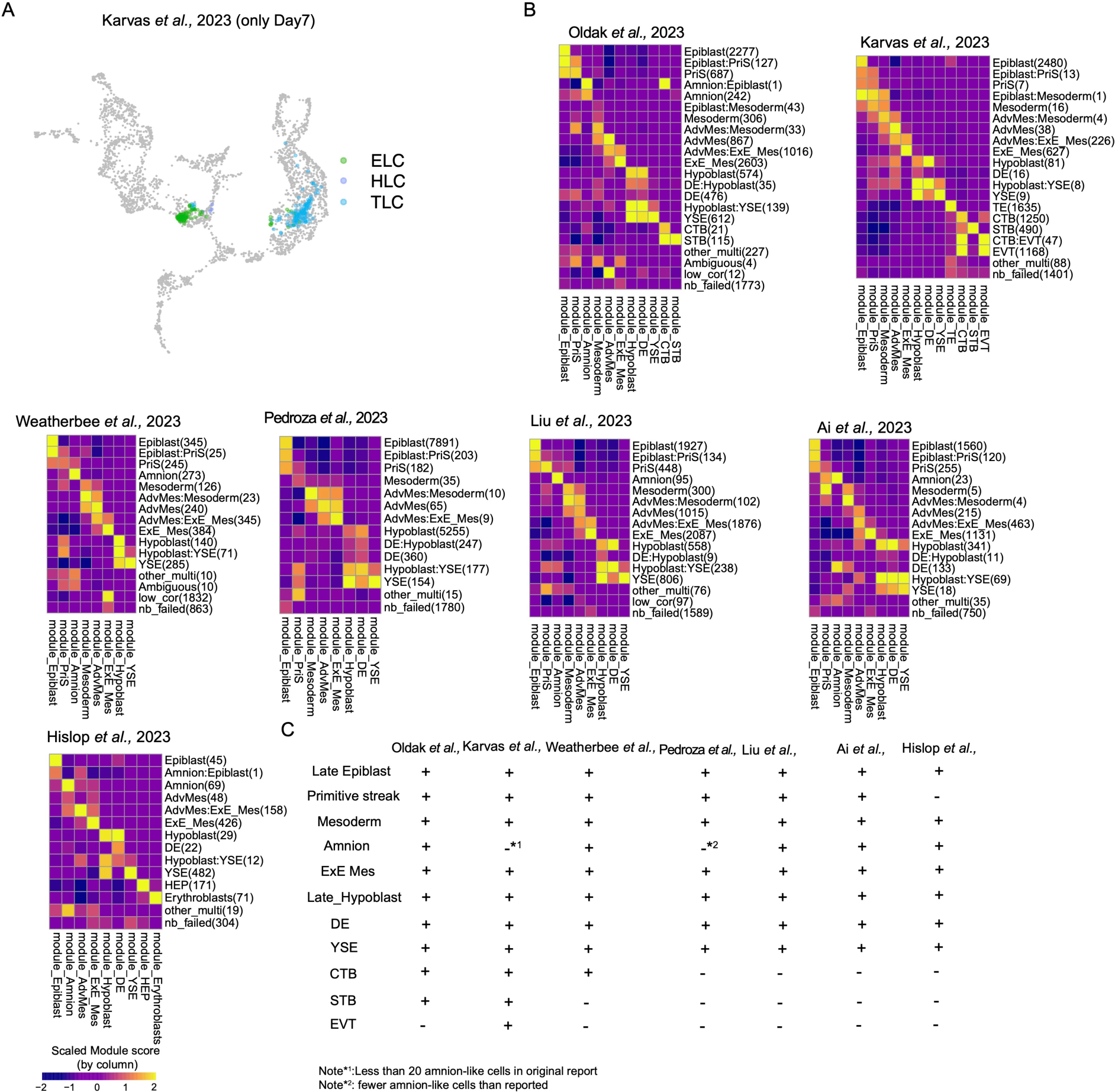
(A) Highlighted cells from the Day 7 blastoids (neighbourhood nodes) from Kavas et al., 2023. The colour of each data point represents the cell annotations retrieved from original publication. (B) Module score of corresponding predicted lineages in seven postimplantation models. (C) Checking the existence of post-implantation lineage-like cells in post-implantation embryo models.

**Figure S13.**
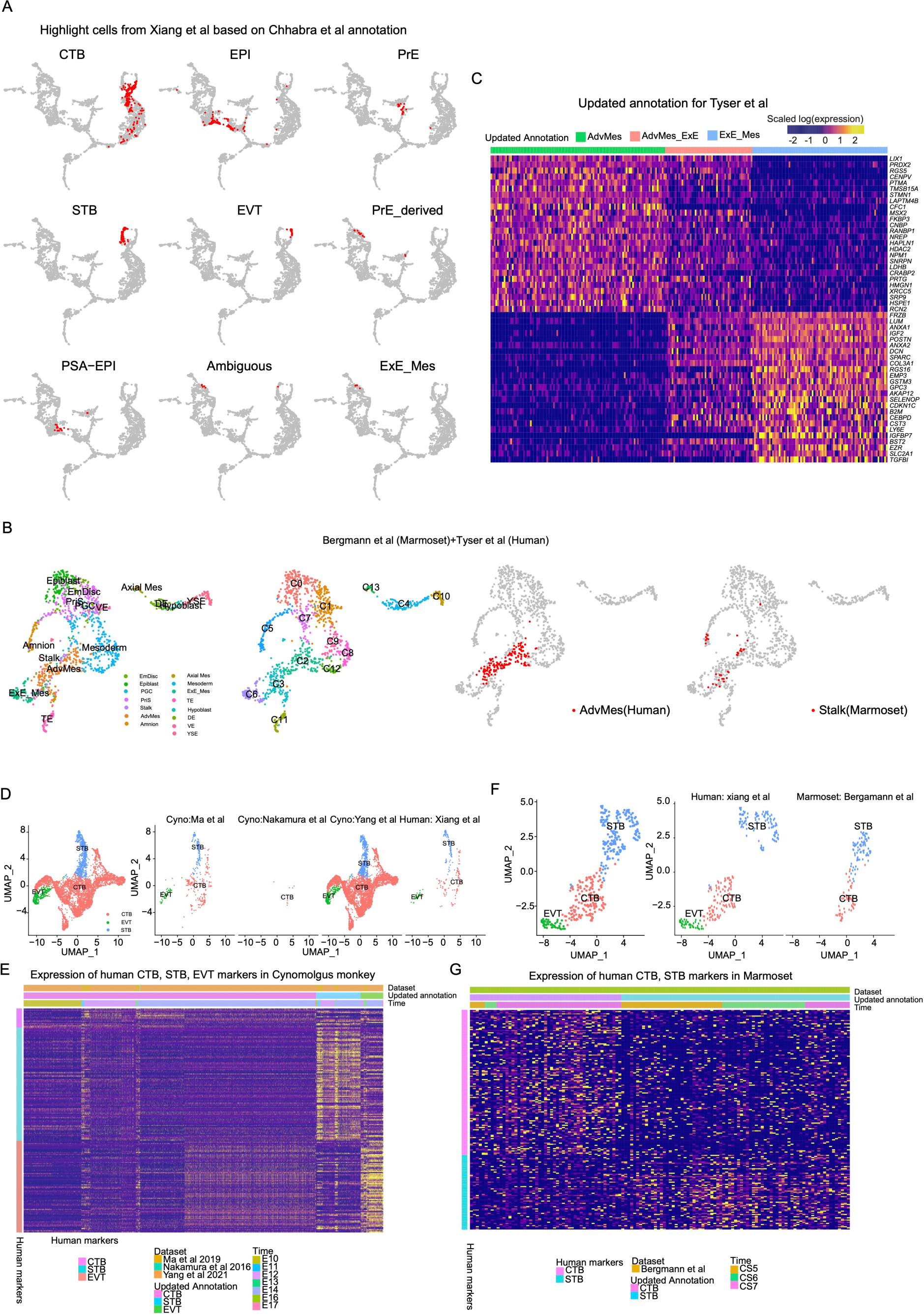
(A) Highlighted cells from Xiang et al. based on the annotation from Chhabra et al. (B) UMAP projection of Mutual Nearest Neighbours (MNN) cross-species integration, including CS7 cells from human (Tyser et al) and the Marmoset embryo (Begamann et al). The stalk cells from Marmoset and the human previously advanced mesoderm were highlighted separately. (C) Heatmap showing DEGs between Advanced Mesoderm (not belonging to cluster C3) and human extraembryonic mesoderm cells in human advanced mesoderm cells, reannotated extraembryonic mesoderm cells (AdvMes_ExE), and human embryo extraembryonic mesoderm cells. (D) UMAP projection of Mutual Nearest Neighbours (MNN) cross-species integration, including human and cynomolgus monkey postimplantation TE cells. (E) Heatmap showing the markers for human CTB, STB, and EVT which were also differentially expressed in identified cynomolgus monkey CTB, STB, and EVT cells.

## Notes

http://petropoulos-lanner-labs.clintec.ki.se/

### Competing Interest Statement

The authors have declared no competing interest.

### Summary of Updates

The manuscript is extensively revised and extended.

